# Metrics for conservation success: using the ‘Bird-Friendliness Index’ to evaluate grassland and aridland bird community resilience across the Northern Great Plains ecosystem

**DOI:** 10.1101/868356

**Authors:** Nicole L. Michel, Curtis Burkhalter, Chad B. Wilsey, Matt Holloran, Alison Holloran, Gary M. Langham

**Affiliations:** National Audubon Society; Operational Conservation; Audubon Rockies

**Keywords:** Grassland birds, aridland birds, conservation, management, bird-friendly, ranching, grazing, working lands, accountable conservation, indices

## Abstract

**Aim:** Evaluating conservation effectiveness is essential to protect at-risk species and to maximize the limited resources available to land managers. Over 60% of North American grassland and aridlands have been lost since the 1800s. Birds in these habitats are among the most imperiled in North America, yet most remaining habitats are unprotected. Despite the need to measure impact, conservation efforts on private and working lands are rarely evaluated, due in part to limited availability of suitable methods.

**Location:** Northern Great Plains

**Methods:** We developed a novel metric to evaluate grassland and aridland bird community response to habitat management practices, the Bird-Friendliness Index (BFI), consisting of density estimates of grassland and aridland birds weighted by conservation need and a functional diversity metric to incorporate resiliency. We used the BFI to inform three assessments: 1) a spatial prioritization to identify ecologically significant areas for grassland and aridland birds, 2) estimation of temporal trends in grassland and aridland bird community resilience, and 3) evaluation of the effects of land management practices on grassland and aridland bird communities.

**Results:** The most resilient bird communities were found in the Prairie Potholes region of Alberta, Saskatchewan, northern Montana, and North Dakota, and the lowest BFI values in the southern and western regions of the Northern Great Plains. BFI values varied little over time on average, but trends varied regionally, largely in response to interannual relative variability in grassland and aridland bird densities.

**Main conclusions:** BFI values increased in response to simulated habitat management, suggesting that practices recommended for use in bird-friendly grassland habitat management plans will increase the abundance and resilience of the grassland and aridland bird community, and will be detected using the BFI. The BFI is a tool by which conservationists and managers can carry out accountable conservation now and into the future.

## 1 INTRODUCTION

Grasslands and aridlands, and the birds that rely on them, are under threat. Over 60% of North America’s grasslands have been lost since the 1800s, with some of the greatest losses in the Northern Great Plains Tallgrass and Mixed-grass Prairie regions (Comer, Hak, Kindscher, Muldavin, & Singhurst, 2018). Though much of the grassland loss occurred in the nineteenth and twentieth centuries, agricultural conversion continues today (Samson, Knopf, & Ostlie, 2004). For example, 3 million acres of cropland – a less-suitable habitat for grassland birds – were created during 2008-2012, 77% of which were converted from grassland (Lark, Salmon, & Gibbs, 2015). Additionally, grassland birds face a variety of ongoing threats including agricultural intensification, pesticides, and energy development (Stanton, Morrissey, & Clark, 2018). Consequently, populations of grassland-dependent birds have declined by an estimated 53.3% since 1966, with 74.2% of North American grassland bird species declining and 27% of species considered to be of high conservation concern (North American Bird Conservation Initiative, 2016; Rosenberg et al., 2019). Moreover, grassland birds with specialized habitats on their breeding and/or wintering ranges have experienced greater population declines than habitat generalists (Correll, Strasser, Green, & Panjabi, 2019). Aridlands similarly face multiple threats, including habitat loss due to urban sprawl and energy development, drought, desertification, and invasive species (Ryan et al., 2008; North American Bird Conservation Initiative, U.S. Committee, 2014). In response, aridland bird species have declined an average of 17-46% since 1968, with 56.5% of species declining and 39% of species of conservation concern (North American Bird Conservation Initiative, U.S. Committee, 2014; North American Bird Conservation Initiative, 2016; Rosenberg et al., 2019).

Private lands offer a potential source of hope for grassland and aridland birds, as 84% of grasslands and 44% of aridlands are held in private landownership (Askins et al., 2007; North American Bird Conservation Initiative, U.S. Committee, 2011). Over 90% of the breeding distribution of seven obligate grassland-breeding birds is on private lands, including the Eastern Meadowlark (*Sturnella magna*) and Dickcissel (*Spiza americana*; North American Bird Conservation Initiative, U.S. Committee, 2013). As a result, private lands are essential to ensure conservation of imperiled grassland and aridland birds in the future (North American Bird Conservation Initiative, 2016).

Organizations and managers that seek to conserve imperiled birds and their habitats face a variety of challenges, not the least of which is optimizing the use of limited funds and allocating them in demonstrably successful ways. Ensuring conservation success requires identifying solid goals and objectives, prioritizing lands for conservation, taking appropriate conservation actions in priority locations to meet these goals, and identifying a means by which to measure the results of conservation actions (Pullin, Sutherland, Gardner, Kapos, & Fa, 2013). These efforts form an adaptive management cycle in which actions are undertaken, responses are evaluated, and consequently actions are revised (Nichols & Williams, 2006). Yet prioritizing lands for conservation of entire communities and quantifying the success of conservation efforts are challenging and often elusive endeavors due to, e.g., species-specific responses to conservation actions, and the spatial and temporal disconnects between action and response (Kapos et al., 2008, 2009). Metrics that effectively capture the status of ecological communities, community change over time, and the impacts of conservation actions on these communities are needed to support effective and accountable conservation.

Conservation metrics often rely on evaluations of habitat quality for single species that are treated as indicators (Simberloff, 1998). Yet single species rarely serve as effective proxies for larger communities, due to limited overlap in habitat requirements, functional roles, and responses to management actions among species (Cushman, McKelvey, Noon, & McGarigal, 2010; Johnson, Igl, Shaffer, & DeLong, 2019; Larsen, Bladt, & Rahbek, 2009; Lindenmayer, Barton, & Pierson, 2015; Winter, Johnson, & Shaffer, 2005). Single-species indicator approaches are particularly ill-suited to communities with threatened species, as more common species often perform poorly as surrogates for species of conservation concern (Stephens, Dinger, & Alexander, 2019). Indeed, in many cases surrogates do not exist or poorly predict responses of imperiled species (Lindenmayer et al., 2015; Silvano, Guyer, Steury, & Grand, 2017). Additionally, species-specific abundance alone is rarely a reliable predictor of habitat quality (Johnson, 2008). Instead, metrics incorporating distributions, habitat relationships, and diversity of multiple species better encapsulate the response of ecological communities to conservation actions (Goyert et al., 2016; Nuttle, Leidolf, Burger, & Loiselle, 2003).

By incorporating suites of species using habitat in different ways in multi-species metrics, we can incorporate the variability among species in their habitat use, foraging preferences, and functional contributions to ecosystems (e.g., pest control, seed dispersal; Şekercioğlu, Wenny, & Whelan, 2016). Together, these suites of species and their associated functional traits and services influence the integrity (i.e., ecosystem structure, composition and function; Karr, 1981) and resilience (i.e., ability to resist or recover from disturbances; Scheffer, Carpenter, Dakos, & van Nes, 2015) of the larger ecological community (Fischer et al., 2007). Functional traits such as diet, body size and habitat use influence species’ ecological impacts, and communities composed of species with complementary suites of traits will have greater ecological integrity and resilience (Cardinale et al., 2012; McGill, Enquist, Weiher, & Westoby, 2006). Consequently, functional diversity measures incorporating functional traits and species abundance provide an indirect way to measure resilience and integrity (Standish et al., 2014), and as such are increasingly used in large-scale assessments of North American bird and ecological communities (Schipper et al., 2016; Schleuter, Daufresne, Massol, & Argillier, 2010).

In order to evaluate the success of habitat enhancement in conserving birds and their habitats on private lands in North American grasslands and aridlands, we developed a metric evaluating bird community response to habitat management practices. This metric, the Bird-Friendliness Index (BFI), uses avian count, functional trait, and conservation status data together with remotely sensed environmental data to evaluate the capacity of a landscape to support an abundant, diverse, and resilient bird community. The BFI was designed to enable inference at multiple spatial and temporal scales, and incorporates standardization methods that facilitate spatial comparisons among North American grasslands and aridlands spanning expansive climatic and community gradients. Here we demonstrate the use of the BFI to inform three assessments: 1) spatial prioritization to identify ecologically significant areas for grassland and aridland birds, 2) estimation of temporal trends in relative grassland and aridland bird community resilience, and 3) evaluation of the effects of land management practices on grassland bird communities. We use the Northern Great Plains of North America as a case study, and highlight the insights the BFI provides into the plight of grassland and aridland birds and the integrity and resilience of the larger ecosystems in which they reside.

## 2 METHODS

### 2.1 Bird-Friendliness Index overview

The BFI takes into account species density, conservation status, and diversity of the entire community of grassland birds at a site. Bird abundance, and changes in relative abundance over time, are presently the most common tool for monitoring and evaluating landbird abundance (e.g., Rosenberg et al., 2016; Sauer et al., 2017), and form the foundation of the BFI. Including conservation status ensures that the BFI does not underestimate conservation need by diluting the contribution of rare species (Beissinger, 2000). Lastly, the BFI also includes a measure of functional diversity as a measure of the intactness and resilience of both the grassland and aridland bird community and their larger ecological community (Fischer et al., 2007).

The BFI is the sum product of estimated avian density and conservation status times a measure of functional diversity based on species traits (i.e., diet, foraging strata, and body mass), as follows:

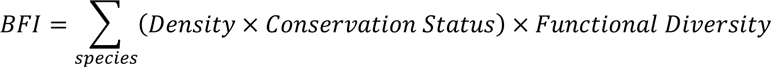

It is calculated in six steps, all of which are calculated at the resolution of 1 km^2^ grid cells. This resolution was chosen because it represents the smallest scale at which the avian survey data used here are summarized (see *Avian data*), and it is large enough to contain entire home ranges of most study species. First, densities for each species were estimated for surveyed grid cells using point counts that include one or more auxiliary methods enabling estimation of detection-corrected density (e.g., distance or time of first detection; Matsuoka et al., 2014). Second, individual species models relating density to remotely-sensed environmental covariates were built and used to predict densities across the study area. Third, species densities were multiplied by a conservation score for each species, so that threatened and rare species carry a greater weight. Fourth, conservation status-corrected densities for each species were summed within each grid cell. Fifth, summed conservation-corrected densities for each grid cell were multiplied by an index of functional diversity calculated using the predicted densities (Step 2) and corresponding functional traits for the species present within a grid cell. Finally, BFI values were standardized to enable spatial comparison by scaling from zero to one using a logistic transformation across the study area, Bird Conservation Region (BCR; Bird Studies Canada and North American Bird Conservation Initiative, 2014), or other region of interest. BFI values can be easily aggregated to larger scales (e.g., ranch, state, BCR) by calculating the mean or median of grid cells included within the area of interest.

The standardization process is a critical component of the BFI. Standardization expresses each cell’s bird-friendliness relative to the study area, producing a ranked index scaled from zero to one that is easily interpreted. Furthermore, it enables comparisons of BFI values from differing grassland or aridland types and climatic zones across North America. Finally, standardizing relative to a larger geographic region improves tracking of local-scale management effects by controlling for interannual variation due to large-scale processes, e.g. weather, density-dependent cycles, and background population-level trends.

### 2.2 Study area

Our study area encompasses the Northern Great Plains ecosystem, as defined by the West-Central Semi-arid Prairies level II ecoregion (Commission for Environmental Cooperation (CEC), 2009). This region stretches from the Prairie Pothole region of south-central Canada to Nebraska and from eastern North Dakota to western Montana (Figure 1). It consists primarily of Northwestern Great Plains Mixed-grass Prairie, with some Northern Great Plains Fescue Mixed-grass Prairie along the northern edge of the study area, and Western Great Plains Shortgrass Prairie and Western Great Plains Sand Prairie to the south (Comer et al., 2018). The study area also includes extensive arid sagebrush steppe habitat intermixed with grasslands, ranging from 1-25% of regional land cover in Montana and the Dakotas, to >75% cover in northeastern Wyoming (Connelly, Knick, Schroeder, & Stiver, 2004).

**FIGURE 1.**
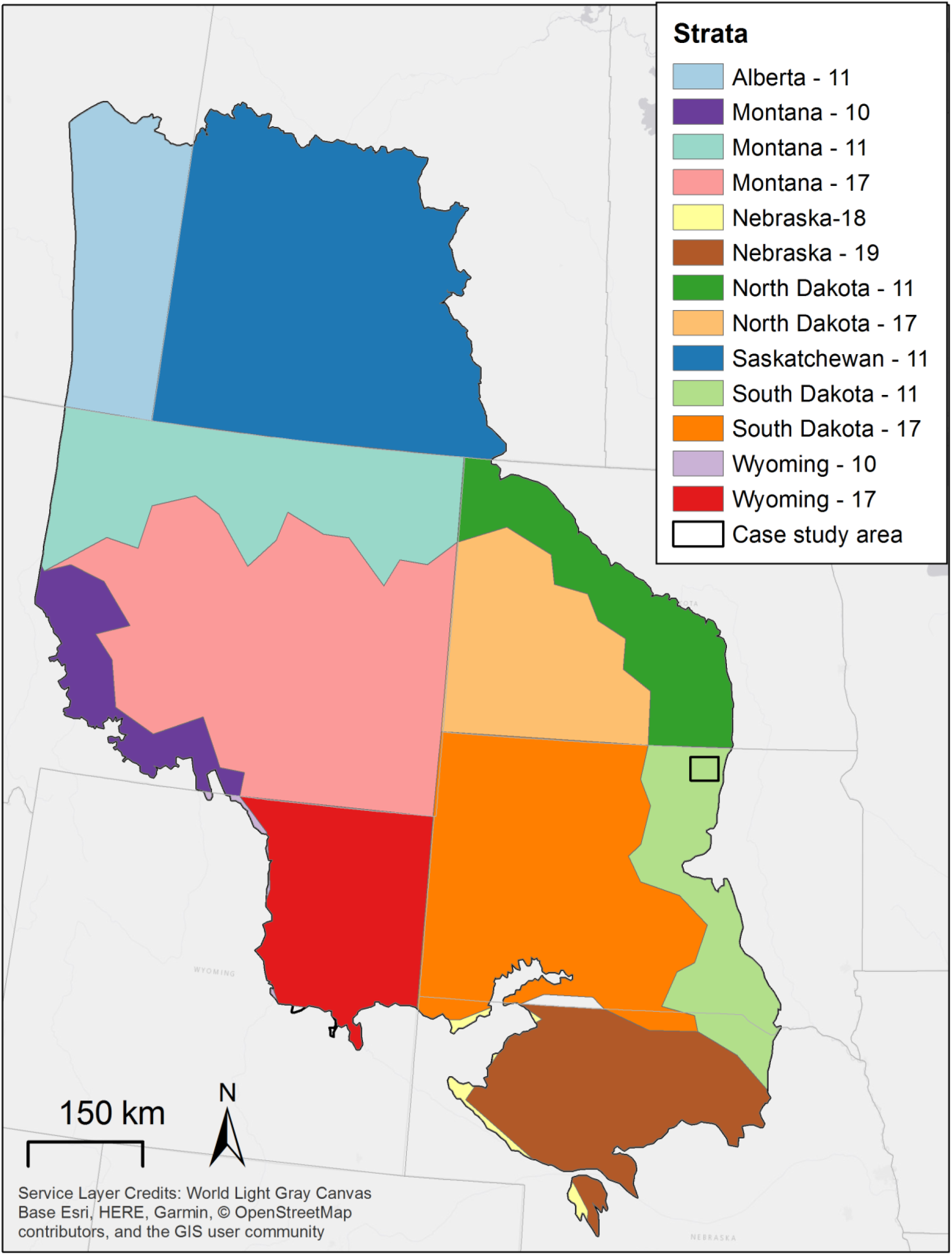
Map of Northern Great Plains study area showing boundaries of strata, defined as unique combinations of states/provinces and Bird Conservation Regions (for BCR names see *Methods*), and location of the management case study.

### 2.3 Study species

We used the 2016 State of North America’s Birds species assessment to identify 107 bird species that use grasslands or aridlands as their primary or secondary breeding habitat (North American Bird Conservation Initiative, 2016). We selected grassland and aridland birds because the two habitats are intermixed at relatively fine spatial scales in this region, and private landholdings on which management practices are implemented often include both habitat types and, consequently, communities. This list formed the pool of species evaluated for inclusion in the BFI contingent on data availability in the study area (see *Avian data*).

### 2.4 Avian data

We used avian point count data collected under Bird Conservancy of the Rockies’s Integrated Bird Monitoring in Bird Conservation Regions (IMBCR) program (Woiderski et al., 2018). The IMBCR uses a hierarchically nested sampling design in which spatially balanced random stratification is used to select 1 km^2^ grid cells within BCRs. Each grid cell includes up to 16 point count locations evenly spaced at distances of 250 m. At each point, five- (2009) or six-minute (2010-2014) unlimited-distance point count surveys were conducted between one-half hour before and five hours after sunrise during the late spring and early summer. Observers recorded species, age (where possible), first minute of observation, type of detection (call, song, visual), day, time of day (2010-2014 only), and primary habitat type at the point, and measured distance to all individuals and flocks using a laser rangefinder. We excluded flyover observations, juvenile birds, and observations with missing minutes or distances.

We assigned IMBCR survey points to spatial strata defined as unique combinations of BCRs and states or provinces, as used in hierarchical modeling of Audubon Christmas Bird Count data (Figure 1; Soykan et al., 2016). We used a subset of the IMBCR data for strata in which sampling occurred annually from 2009 to 2014. This limited us to surveys conducted within Montana, North Dakota, South Dakota, Wyoming and Nebraska. Our study area encompassed five BCRs: Northern Rockies (BCR 10), Prairie Potholes (BCR 11), Badlands and Prairies (BCR 17), Shortgrass Prairie (BCR 18), and Central Mixed Grass Prairie (BCR 19).

Our dataset spanned 2009-2014 and included a total of 1,102 unique 1 km^2^ grid cells, with 242 – 594 grid cells surveyed annually and an average of 11 points sampled per grid cell. Of these, 34 species had sufficient data from which to estimate density (see *Density estimation*) and were included in the BFI (Table 1). These included 26 species that use grassland habitats and 12 species that use aridlands as a major breeding habitat. Four species (Bell’s Vireo, Lark Sparrow, Loggerhead Shrike, and Western Kingbird) use both grasslands and aridlands as major breeding habitats (Table 1), and many species use both habitats facultatively. The final dataset for these 34 species included 45,152 observations of 99,842 birds.

**TABLE 1.** Grassland and aridland bird species included in Bird-Friendliness Index estimation for the NGP, their major breeding habitats, functional species grouping, breeding season Combined Conservation Score (CCSb), and mean density (birds/km^2^) at surveyed grid cells. Conservation scores and habitat associations come from the 2016 State of the Birds (North American Bird Conservation Initiative, 2016).

| Common Name | Scientific Name | Major Breeding Habitats | Functional Species | CCSb | Density (mean ± SE) |
| --- | --- | --- | --- | --- | --- |
| Baird's Sparrow | *Ammodramus bairdii* | Grassland | Ground-foraging invert- & granivores | 15 | 1.69 ± 0.16 |
| Bell's Vireo | *Vireo bellii* | Aridland, forest | Mid- to upperstory foraging invertivores | 11 | 0.08 ± 0.02 |
| Bobolink | *Dolichonyx oryzivorus* | Grassland | Ground-foraging invert- & frugivores | 12 | 3.53 ± 0.30 |
| Brewer's Sparrow | *Spizella breweri* | Aridland | Ground-foraging invert- & granivores | 11 | 7.46 ± 0.48 |
| Burrowing Owl | *Athene cunicularia* | Grassland, aridland | Large ground-foraging invert- & vertivores | 12 | 0.02 ± 0.006 |
| Canyon Wren | *Catherpes mexicanus* | Aridland | Ground- or aerial-foraging invertivores | 9 | 0.02 ± 0.004 |
| Chestnut-collared Longspur | *Calcarius ornatus* | Grassland | Ground-foraging invert- & granivores | 15 | 10.96 ± 0.81 |
| Clay-colored Sparrow | *Spizella pallida* | Grassland | Ground-foraging invert- & granivores | 9 | 2.61 ± 0.22 |
| Dickcissel | *Spiza americana* | Grassland | Ground-foraging invert- & granivores | 10 | 0.60 ± 0.12 |
| Eastern Kingbird | *Tyrannus tyrannus* | Grassland, forest | Understory-foraging invert-, frug- & granivores | 10 | 3.12 ± 0.15 |
| Ferruginous Hawk | *Buteo regalis* | Grassland | Large ground-foraging vertivores | 10 | 0.005 ± 0.001 |
| Grasshopper Sparrow | *Ammodramus savannarum* | Grassland | Ground- or aerial-foraging invertivores | 12 | 45.28 ± 1.48 |
| Greater Prairie-chicken | *Tympanuchus cupido* | Grassland | Ground-foraging invert- & granivores | 16 | 0.006 ± 0.001 |
| Greater Sage-grouse | *Centrocercus urophasianus* | Aridland | Large ground-foraging herbivores | 15 | 0.01 ± 0.003 |
| Green-tailed Towhee | *Pipilo chlorurus* | Aridland | Ground-foraging invert- & frugivores | 11 | 0.23 ± 0.06 |
| Horned Lark | *Eremophila alpestris* | Grassland, tundra | Ground-foraging invert- & granivores | 9 | 22.84 ± 0.94 |
| Lark Bunting | *Calamospiza melanocorys* | Grassland | Ground-foraging invert- & granivores | 12 | 17.98 ± 0.92 |
| Lark Sparrow | *Chondestes grammacus* | Grassland, aridland | Ground-foraging invert- & granivores | 9 | 11.63 ± 0.47 |
| Loggerhead Shrike | *Lanius ludovicianus* | Grassland, aridland | Small ground-foraging invert-,<br>vertivores & scavengers | 11 | 0.45 ± 0.04 |
| Long-billed Curlew | *Numenius americanus* | Grassland | Ground- or aerial-foraging invertivores | 14 | 0.22 ± 0.03 |
| McCown's Longspur | *Rhynchophanes mccownii* | Grassland | Ground-foraging invert- & granivores | 15 | 0.84 ± 0.14 |
| Mountain Plover | *Charadrius montanus* | Grassland, wetland | Ground- or aerial-foraging invertivores | 15 | 0.05 ± 0.01 |
| Northern Harrier | *Circus cyaneus* | Grassland, wetland | Large ground-foraging invert-,<br>vertivores & scavengers | 11 | 0.39 ± 0.02 |
| Rock Wren | *Salpinctes obsoletus* | Aridland | Small ground-foraging invert- &<br>vertivores | 10 | 1.39 ± 0.08 |
| Sage Thrasher | *Oreoscoptes montanus* | Aridland | Ground- or aerial-foraging invertivores | 11 | 0.08 ± 0.01 |
| Savannah Sparrow | *Passerculus sandwichensis* | Grassland, tundra | Ground-foraging invert- & frugivores | 8 | 2.19 ± 0.19 |
| Sharp-tailed Grouse | *Tympanuchus phasianellus* | Grassland, forest | Midsized ground-foraging herb- &<br>granivores | 10 | 0.17 ± 0.02 |
| Sprague's Pipit | *Anthus spragueii* | Grassland | Ground- or aerial-foraging invertivores | 14 | 0.77 ± 0.07 |
| Swainson's Hawk | *Buteo swainsoni* | Grassland | Large ground-foraging invert- & vertivores | 9 | 0.04 ± 0.003 |
| Upland Sandpiper | *Bartramia longicauda* | Grassland | Ground- or aerial-foraging invertivores | 10 | 1.50 ± 0.07 |
| Vesper Sparrow | *Pooecetes gramineus* | Grassland | Ground-foraging invert- & granivores | 10 | 9.85 ± 0.35 |
| Western Kingbird | *Tyrannus verticalis* | Grassland, aridland | Mid- to upperstory foraging invertivores | 7 | 2.36 ± 0.13 |
| Western Meadowlark | *Sturnella neglecta* | Grassland | Ground-foraging invert- & granivores | 9 | 28.38 ± 0.58 |
| White-throated Swift | *Aeronautes saxatalis* | Aridland, forest | Ground- or aerial-foraging invertivores | 11 | 0.43 ± 0.10 |

### 2.5 Environmental data

We used 17 environmental variables in individual species models to predict avian densities across the study area (see *Density prediction*; Table S1). All environmental variables were sampled for 799,015 grid cells of 1 km^2^, covering the study area and aligning with the avian density grid cells. We used statistically downscaled climate normals from 1981-2010 derived from the Climatic Research Unit Timeseries 3.22 dataset to represent long-term climatic conditions (Wang, Hamann, Spittlehouse, & Carroll, 2016). We obtained annually-derived ecosystem metrics (e.g., net primary productivity, soil moisture, evaporation) from the NASA CASA terrestrial ecosystem model (Potter et al., 1993; Potter, Klooster, Huete, & Genovese, 2007). We calculated proportion cover (cropland, grassland, shrubland, wetland), mean patch size and patch cohesion index (a measure of physical connectedness; McGarigal, Cushman, Neel, & Ene, 2002; grassland only) from the Commission for Environmental Cooperation’s (CEC) North American Environmental Atlas 2010 landcover dataset (Canada Centre for Remote Sensing (CCRS) et al., 2017). All landcover metrics were calculated using package *SDMTools* (VanDerWal, Falconi, Januchowski, Shoo, & Storlie, 2014) in R version 3.5.2 (R Core Team, 2018). Terrain ruggedness was derived from a digital elevation model to quantify topographic heterogeneity (Riley, Degloria, & Elliot, 1999). Environmental variables were selected based on known relationships with distribution and abundance of grassland and aridland birds (e.g., Best et al., 1997; Fedy, Devries, Howerter, & Row, 2018; Fisher & Davis, 2010; George, Fowler, Knight, & McEwen, 1992; Harrower et al., 2017; Niemuth, Solberg, & Shaffer, 2008; Pool, Panjabi, Macias-Duarte, & Solhjem, 2014; Renfrew et al., 2013; Wiens, 1974).

### 2.6 Density estimation

As the avian dataset included both time of first detection and distance of observations, we used a formulation of time removal and distance sampling models developed by Sólymos et al., (2013) and implemented in R package *detect* (Sólymos, Moreno, & Lele, 2016). We estimated point-level density corrected by two components of detection probability, availability and perceptibility, using conditional multinomial maximum likelihood estimation. Availability – the probability that a bird provided a visual or auditory cue during sampling and was thus available to be detected – was estimated using the minute of first observation. Each individual was counted only once, thus individuals were ‘removed’ once detected. Perceptibility – the conditional probability that birds available for detection were actually detected – was estimated as a function of distance from observer (Sólymos et al., 2013).

We combined data from 2009-2014 in a single model per species to maximize the number of species with sufficient data for modeling, but allowed availability and perceptibility to vary among years. We obtained model convergence for species with 23 or more observations during all years. The number of observations per species ranged from 23 (Canyon Wren) to 34,857 (Western Meadowlark), with a median value of 544 ± 6,533 SD individuals. To account for species absence or non-detection, we created zero observation records for all surveys at which each species was not observed.

We grouped observations into three time periods: 0-2 minute, 2-4 minute, 4-6 minute (4-5 minutes in 2009), and three distance bins: 0-50 m, 51-100 m, and >100 m. Availability was estimated as the singing rate, defined as the probability a bird sang or gave a visual cue during a survey. Singing rate was collapsed across distance bins and modeled as a log-linear function. We controlled for temporal and environmental variables affecting singing rate by including linear terms for year, primary habitat, and spatial strata, and both linear and quadratic terms for day of year, in addition to the intercept. We calculated perceptibility using the effective detection radius, defined as the distance beyond which as many birds were detected as remained undetected within. We controlled for temporal and environmental variables affecting perceptibility by including linear terms for year, primary habitat, spatial strata, and surveyor identity, and both linear and quadratic terms for day of year, in addition to the intercept. Models including all combinations of covariates, and no covariates, were built and model selection was conducted using AIC (Burnham & Anderson, 2002).

Singing rate and effective area sampled (in ha), defined as the area within the effective detection radius, were calculated at the point level then used to estimate a correction factor for each survey accounting for imperfect detection. Correction factors were log transformed and entered as offsets in generalized linear mixed models calculated using R package *lme4* (Bates, Mächler, Bolker, & Walker, 2015). We modeled detection-corrected annual density at the point level using a Poisson distribution and included primary habitat as a fixed effect, and grid cell and year as random effects. We left-truncated density estimates at 0.0008 birds/ha, equivalent to the density estimate for a point with a single bird observed at the maximum observed distance in the dataset (2000 m). Annual densities at the 1 km^2^ grid cell level were estimated by taking the median of annual point-level densities (birds/ha) within each surveyed grid cell multiplied by 100.

### 2.7 Species habitat modeling

We built species habitat models, with density as the response variable, for each species using boosted regression trees (BRTs), a machine learning approach that is ideal for modeling complex species-environment relationships with multiple, and often highly correlated, predictors (Elith, Leathwick, & Hastie, 2008). As many species had skewed density distributions with absences in many grid cells, we implemented a hurdle model approach in which we separately modeled occurrence and density. The response variables were presence/absence for the presence model, and detection-corrected abundance scaled to 1 km^2^ for the density model. For the density model, we first rounded densities and used a Poisson distribution. If these models failed to converge, we log-transformed estimated densities to improve fit and used a Gaussian distribution. The predictor variables included the 17 environmental variables described above, plus year (Table S1). Models were fit using R packages *dismo* (Hijmans, Phillips, Leathwick, & Elith, 2015) and *gbm* (Ridgeway, 2017).

BRTs use three regularization parameters to shrink the number of terms in the final model and thus avoid overfitting. Learning rate shrinks the contribution of each tree to the full model, bag fraction specifies the proportion of data selected from the training set at each iteration, and tree complexity sets the number of nodes and, thus, level of interactions between predictors (Elith et al., 2008). We iteratively tuned these parameters to optimize model fit while ensuring a minimum of 1,000 trees using default parameter ranges recommended by Elith et al., (2008): learning rate 1.0*e*^−10^–0.1, bag fraction 0.5–0.75, and tree complexity 1–3. At each step we used 10-fold cross-validated area under the curve (AUC; occurrence model only) and residual deviance (both models) to select the optimal parameter value.

Species distribution models such as these are susceptible to spatial autocorrelation, which can result in biased presence/density-environment relationships (Boria, Olson, Goodman, & Anderson, 2014). We took two steps to reduce residual spatial autocorrelation in the model. First, we conducted geographic filtering by overlaying a 10 km^2^ grid and randomly selecting only one grid cell from each 10 km^2^ grid for inclusion in the analyses (Boria, Olson, Goodman, & Anderson, 2014). We chose this scale based on Wilsey et al., (2019), which tested three geographic resolutions for filtering and found that 10 km produced the greatest model performance. Second, we used spatially stratified cross-validation by dividing the data sets into 12 gridded bins by latitude and longitude, and withholding one latitudinal bin for testing at each fold (Roberts et al., 2017). We tested for residual spatial autocorrelation in the final models using Moran’s I, calculated in R package *ape* (Paradis, Claude, & Strimmer, 2004), and evaluated model performance using cross-validated AUC and TSS (presence only), deviance explained, and correlation. We evaluated the influence of each predictor variable by calculating mean relative importance values for all 34 species.

Because of the randomness introduced by the geographic filtering, as well as the iterative BRT estimation process itself, we produced 25 bootstrapped geographically filtered datasets. We selected 25 bootstrap iterations as there was a maximum of 24 surveyed grid cells within a single 10 km^2^ filtering grid. Each of the 25 presence/absence and density models were used to generate predicted probability of occurrence or density across the entire study area using the 18 predictor variables. For each species, we calculated median predicted probability of occurrence and density for each grid cell and across the 25 bootstraps. We calculated a minimum probability of occurrence threshold using maximum sensitivity + specificity in R package *SDMTools* (VanDerWal et al., 2014), and used this to mask median predicted density such that densities were retained only at grid cells that surpassed the threshold hurdle.

### 2.8 Conservation scores

We incorporated the breeding season Combined Conservation Score from the State of North America’s Birds (North American Bird Conservation Initiative 2016), as these scores were calculated consistently for all native bird species across North America trends using a process developed over many years and updated in response to peer-review. Combined Conservation Sscores for ranged from 7 – 20 based on population size, distribution, threats and trends (Panjabi, Blancher, Dettmers, & Rosenberg, 2012). Species-specific breeding season Combined Conservation Scores were multiplied by predicted densities for each 1 km^2^ grid cell to produce conservation-corrected densities, which were then summed across all 34 species per grid cell. We explored scaling the Combined Conservation Scores by dividing each score by the root mean square (range: 0.60 – 1.37). BFI values calculated with scaled Combined Conservation Scores were directly correlated with BFI values calculated with raw Combined Conservation Scores in each year (ρ = 0.98 – 0.99, *p* < 0.0001). Therefore, we used the raw scores in BFI estimation.

### 2.9 Functional Diversity

Functional diversity indices take a variety of forms, differing in the resolution of the trait information they include and the use of presence versus abundance data (Schipper et al., 2016; Schleuter et al., 2010). We implemented a two-step approach that used environmental filtering to group species into functional groups (Table 1) using both high-resolution continuous traits (i.e., body mass, proportion dietary composition) and discrete categorical traits (i.e., foraging strata) obtained from EltonTraits 1.0 (Wilman et al., 2014). We converted foraging strata from categorical to integer form, subdivided each functional trait into two bins based upon the distribution of values for that trait, and constructed a series of functional groups (Wilsey et al. 2019). These functional groups were then used in place of species identity to calculate a functional Shannon’s diversity index, as both abundance and evenness have been shown to influence ecosystem level processes independent of richness (Petchey & Gaston, 2006).

### 2.10 BFI calculation and standardization

BFI values were calculated by multiplying the summed conservation-weighted densities by functional diversity for each 1 km^2^ grid cell. Raw values were standardized by scaling from zero to one using a logistic distribution, to accommodate the lower limit of zero and the occasional, exceptionally high BFI value. For the spatial prioritization and evaluation of temporal trends we standardized using all grid cells to enable comparison across the study area as a whole. However, for the assessment of bird community response to simulated management we standardized using only grid cell values from the surrounding spatial strata (South Dakota – Badlands and Prairies; Figure 1) to highlight effects of management relative to nearby areas with similar climate and landcover.

### 2.11 Sensitivity analyses

We conducted a sensitivity analysis to assess the relative influence of species density, conservation score, and functional diversity on BFI values. For each of 1,000 bootstrap iterations we sampled each species’ annual densities from a log-normal distribution using estimated means and standard deviations increased by 10% with a total sample size equivalent to the number of 1 km^2^ grid cells (n = 799,015). For each of 1,000 bootstrap samples of conservation scores we drew species values from a uniform distribution with the minimum (4) and maximum (20) conservation weight serving as the bounds of the distribution. For functional diversity we calculated the mean and standard deviation of the functional evenness values across the entire NGPE in 2011 and again increased the standard deviation by 10%. For functional diversity we sampled values 1,000 times from a beta distribution (which produces values ranging from 0-1) using estimated annual means and estimated standard deviations increased by 10%. We calculated a resampled BFI for each iteration, and estimated effect size as the degree of overlap between the distribution of BFI values calculated from the actual data and the distribution of BFI values for each bootstrap sample to determine effect size. Degree of overlap was calculated by using integration of the area of overlap between the two curves. We subtracted the mean overlap from one so that the greatest effect size would correspond to the component with the smallest degree of overlap.

### 2.12 Temporal trends and management case study

We analyzed relative temporal trends in BFI values by spatial strata (i.e., unique combinations of states/provinces and Bird Conservation Regions; Figure 1) and year using general linear models calculated in R package *nlme* (Pinheiro, Bates, DebRoy, Sarkar, & R Core Team, 2018) in R. BFI values were normally distributed, meeting assumptions of Gaussian models. Year (2009-2014), strata, and their interaction were included as fixed effects and strata was included as a random effect in the fully parameterized model. AIC-based model selection was used to select fixed effects for inclusion in the final model. Heterogeneity in variances of strata were accounted for using correlation structures when this term improved the fit of the full model, as determined by AIC. Fixed effects were evaluated using R package *emmeans* (Lenth, 2018) using Tukey adjustments for multiple comparisons, and plotted using *ggplot2* (Wickham, 2016).

We simulated an evaluation of the effects of management actions on a theoretical private property. In these simulations, we modeled the effects of common bird-friendly habitat management actions including reducing cropland cover, CO_2_, evapotranspiration, and N_2_O, and increasing litter biomass, net primary productivity, soil moisture, proportion grassland, grassland cohesion, and grassland patch area. These covariates were selected for the simulation modeling based on known relationships with bird occurrence and abundance, or because they respond to – and therefore serve as proxies for – reduction of cattle rotation rates and consequently grazing intensity, a common recommendation of bird-friendly habitat management plans (Brennan & Kuvlesky, 2005; Johnson et al., 2019; North American Bird Conservation Initiative, 2016; Undersander, Temple, Bartelt, Sample, & Paine, 2000). Grassland cover, patch area, and connectivity; primary productivity; and litter biomass are important predictors of grassland birds, and associated with higher occurrence rates and densities (Fedy et al., 2018; Fisher & Davis, 2010; Harrower et al., 2017; Renfrew et al., 2013). Similarly, many grassland birds have positive associations with soil moisture and negative associations with drought (evapotranspiration), particularly in the drier western Great Plains (George et al., 1992; Niemuth et al., 2008; Wiens, 1974). These environmental variables may be influenced by management practices that increase cover, connectivity, and density of grasses. Conversely, many grassland birds have lower densities in croplands or avoid them altogether (Best et al., 1997; Pool et al., 2014). Cattle grazing increases nitrous oxide (N_2_O) and carbon dioxide (CO_2_) emissions, therefore these variables serve as proxies for cattle grazing intensity (Allard et al., 2007; Oenema, Velthof, Yamulki, & Jarvis, 1997).

Covariate changes were simulated to accumulate incrementally over time at a rate of 10% per year with some random variation included. These simulated environmental conditions were then used to re-calculate bird densities and functional diversity, and re-estimate the BFI each year within that property. We simulated habitat improvement beginning in 2011 to simulate a before-after control-impact design with monitoring for two years prior to management. We then compared changes in simulated BFI values post-management to actual, estimated BFI from 2009 to 2014.

## 3 RESULTS

### 3.1 Bird-Friendliness Index estimation

#### 3.1.1 Estimated density

Densities estimated at surveyed grid cells for the 34 species during 2009-2014 ranged from 0.005 ± 0.001 (Ferruginous Hawk) to 45.28 ± 1.48 (Grasshopper Sparrow; Table 1). The five most abundant species observed across the study area were Grasshopper Sparrow, Western Meadowlark, Horned Lark, Lark Bunting, and Lark Sparrow.

#### 3.1.2 Species habitat modeling

Fit of species habitat models varied among species and data type (presence/absence versus density), but the predictors generally explained 30-40% of the variation in the data after cross-validation using geographic block partitioning. For presence/absence models, cross-validated AUC averaged 0.74 ± 0.02 SE (range: 0.53-0.99), TSS averaged 0.24 ± 0.03 SE (range: 0.02 – 0.72), correlation averaged 0.30 ± 0.03 (range: 0.00 – 0.80), and deviance explained averaged 0.40 ± 0.03 (range: 0.08 – 0.96; Table S2). For density models, cross-validated correlation averaged 0.22 ± 0.02 (range: −0.10 – 0.58), and deviance explained averaged 0.31 ± 0.04 (range: 0.00 – 0.75; Table S2). All Moran’s I values were ≤ 0.05, indicating that the predictors fully accounted for spatial autocorrelation in the data (Table S2).

**TABLE 2.** Model selection table for AIC-based model selection procedures used to select fixed effects for modeling strata-level trends in BFI. Models including an interaction effect also included main effect terms for variables involved in the interaction. All models included an intercept.

| Fixed effects | AIC | ΔAIC |
| --- | --- | --- |
| Year × Strata | -3521947 | 0 |
| Year + Strata | -3507613 | 14334 |
| Strata | -3507611 | 14336 |
| Year | -1679255 | 1842692 |
| Intercept only | -1679254 | 1842693 |

Climate and landcover variables explained the most variation in presence/absence and density of the grassland and aridland birds studied here. Climatic moisture deficit explained the most variation in occurrence (mean ± SE: 18.51 ± 2.80%), followed by terrain ruggedness index (mean ± SE: 12.39 ± 2.52%) and proportion cropland (mean ± SE: 11.22 ± 2.55%; Figure 2A). Proportion cropland cover explained the most variation in density (mean ± SE: 13.07 ± 3.37%), followed by climatic moisture deficit (mean ± SE: 11.42 ± 3.26%) and terrain ruggedness index (mean ± SE: 11.13 ± 3.24%; Figure 2B). Year explained the least variation in both occurrence and density. Though densities and occurrence frequencies varied among years, environmental predictors that varied across both time and space explained more variation than year alone. Relative variable importance varied among species consistent with species-specific habitat preferences, e.g., White-throated Swift density was explained most by terrain ruggedness, while occurrence and density of grassland birds like Upland Sandpiper were explained most by grassland area, patch area, and cohesion (Figures S1-S34).

**FIGURE 2.**
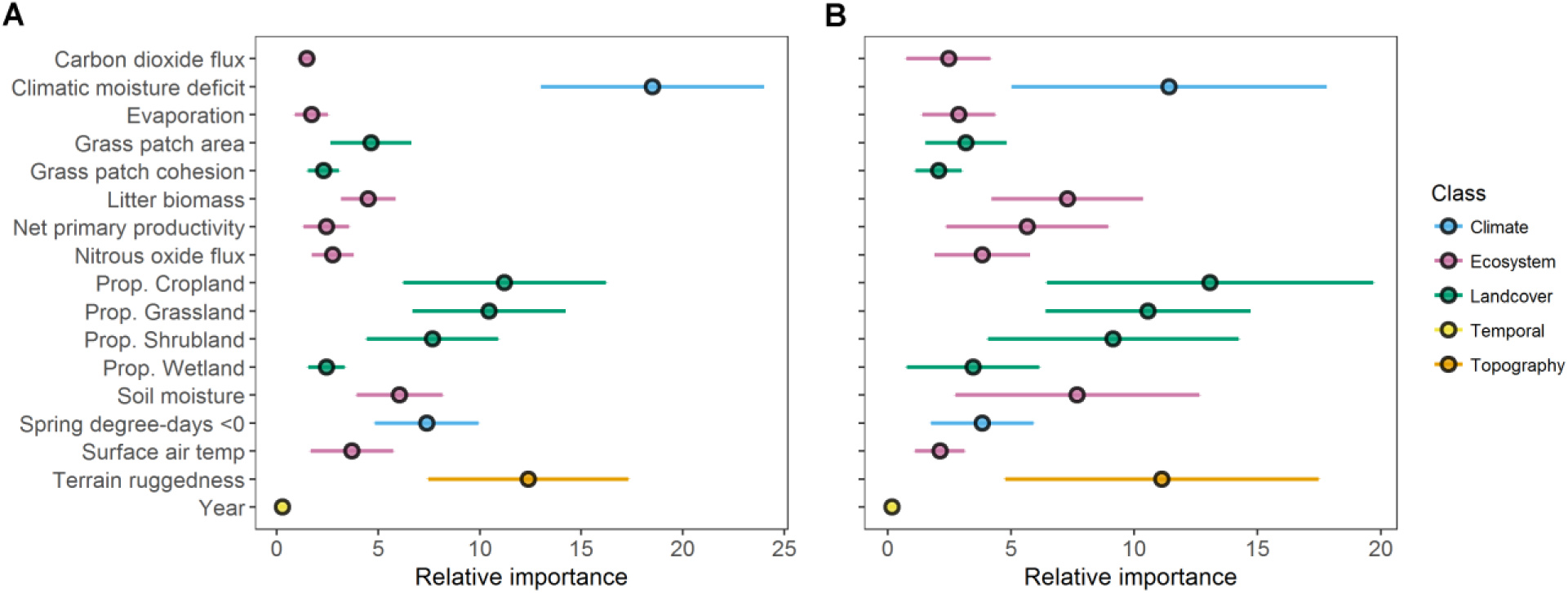
Mean variable importance and 95% confidence intervals for 17 variables used as predictors in presence/absence (A) and density (B) species habitat models for 34 grassland and aridland bird species across the Northern Great Plains during 2009-2014. Climate and landcover variables explained the most variation in occurrence and density.

#### 3.1.3 Functional diversity

Functional diversity averaged 1.34 ± 0.0001 SE (range: 0.00 – 1.99) across the Northern Great Plains during 2009-2014. Functional diversity varied little among years, with the lowest mean diversity of 1.32 ± 0.0003 SE in 2014, and the highest diversity of 1.35 ± 0.0003 SE in 2012.

#### 3.1.4 Bird-Friendliness Index

BFI values showed large- and fine-scale regional variation. The most resilient grassland and aridland bird communities were found in the Prairie Potholes region of Alberta, Saskatchewan, northern Montana, and North Dakota (Figure 3). Conversely, the lowest BFI values were found in the southwestern regions of the study area, notably southern Montana, Wyoming, and southern South Dakota. The regional patterns were generally consistent over time, though high BFI values were more concentrated across the Prairie Potholes in 2013-2014 than earlier years.

**FIGURE 3.**
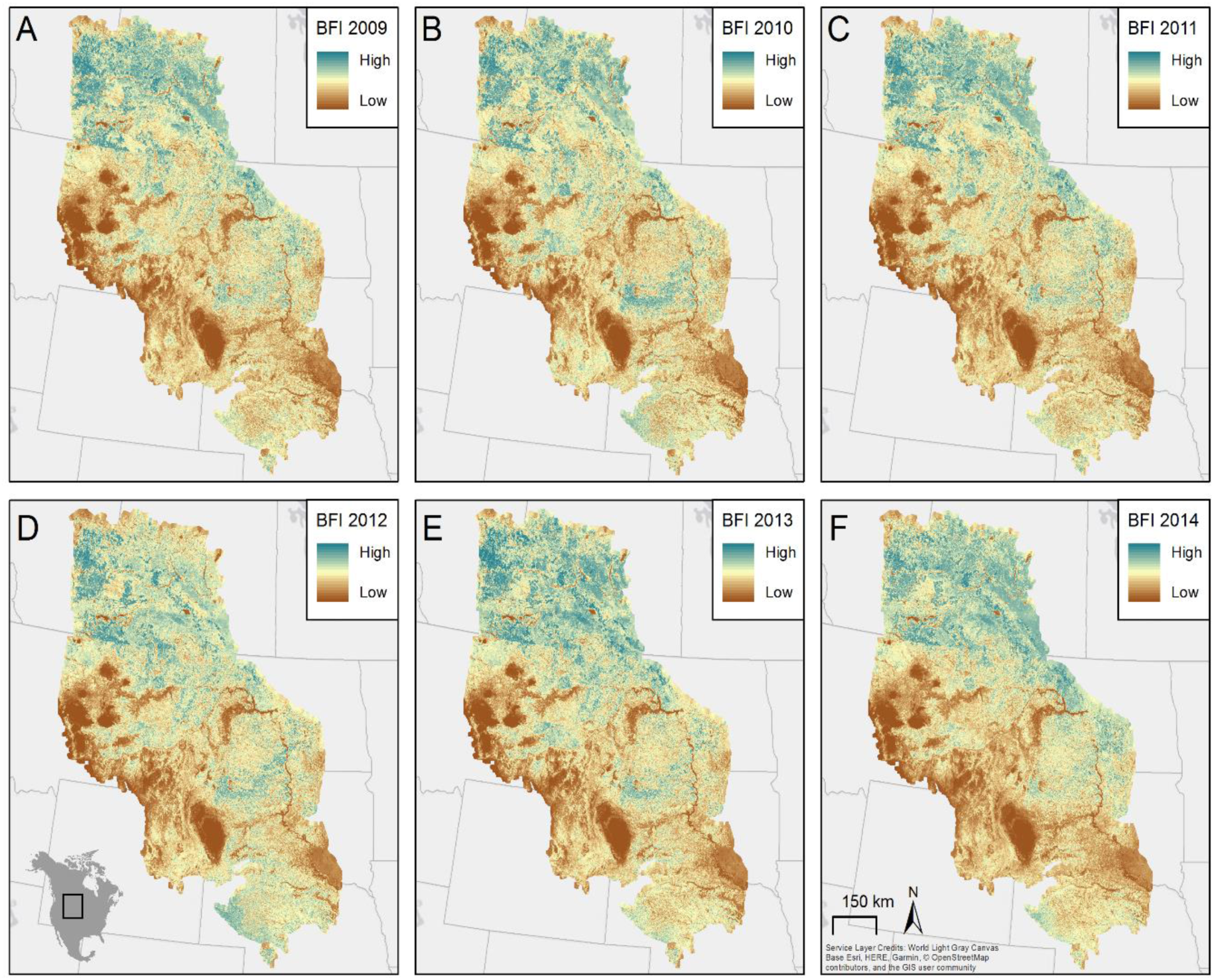
Bird-Friendliness Index across the Northern Great Plains study area for 2009-2014 (A-F). BFI values were highest in Canada, northern Montana, and North Dakota, and lowest in southern Montana, Wyoming, and South Dakota. BFI values showed little variation over time.

#### 3.1.5 Sensitivity analyses

Sensitivity analyses revealed that BFI values were most sensitive to variation in bird densities (Figure 4). BFI calculated with resampled bird densities overlapped with original BFI values by 87.11 ± 0.03%, while resampling conservation scores produced 96.43 ± 0.02% overlap and resampling functional diversity produced 94.41 ± 0.02% overlap (Figure 4).

**FIGURE 4.**
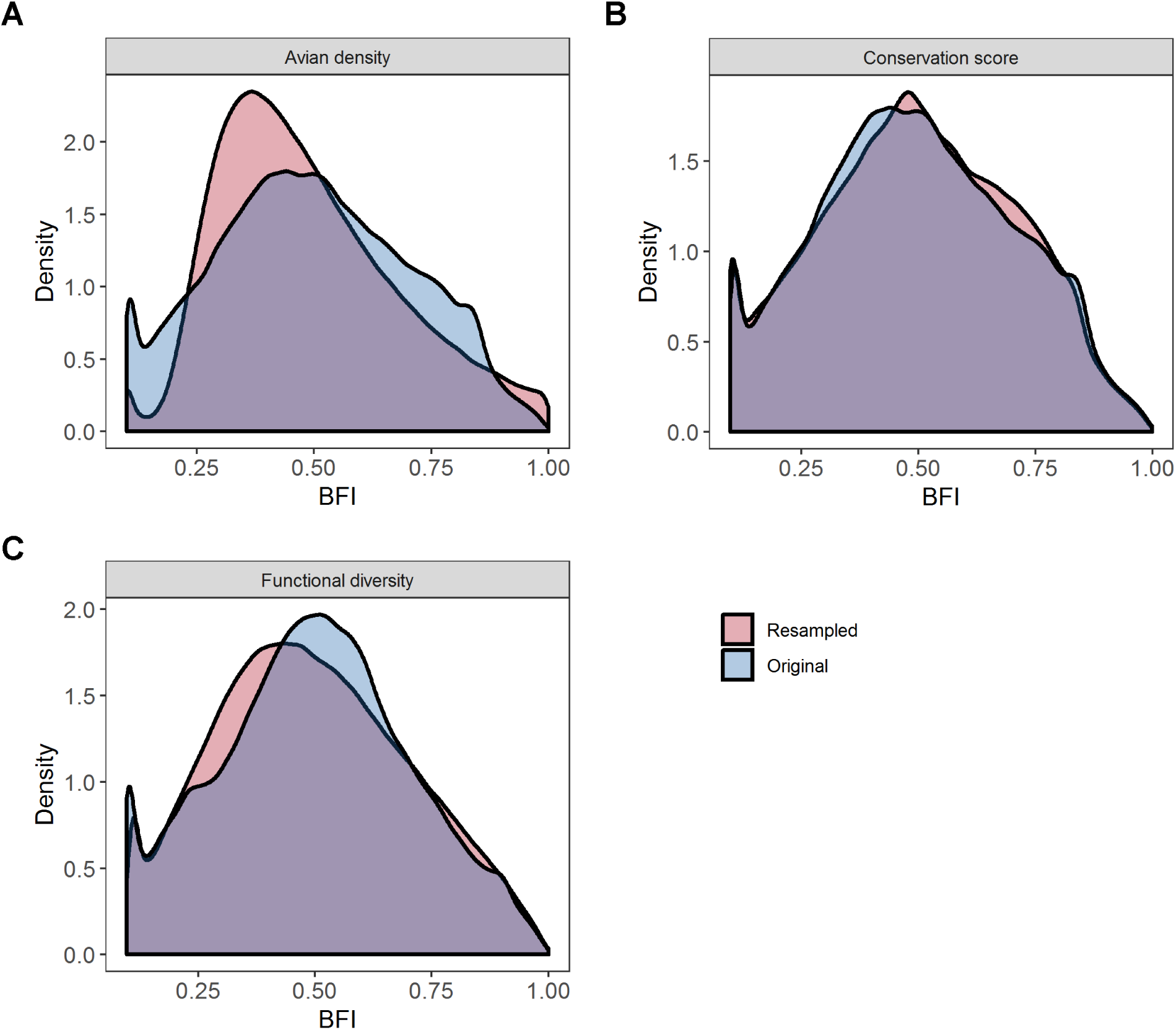
Results of sensitivity analyses evaluating the relative contribution of densities (A), conservation scores (B), and functional diversity (C) to the BFI. The distribution represented by the resampled data is depicted in pink and the distribution represented by the observed data is depicted in blue, with the overlap shown in purple. Variation in densities have the greatest influence on the BFI.

### 3.2 Spatial prioritization

We identified top-priority areas supporting resilient grassland and aridland bird communities as grid cells within the top 20% of BFI values in all years (2009-2014). A total of 427,070 km^2^ fell within the top 20%, concentrated in the Prairie Potholes, North Dakota, northern South Dakota, and central Nebraska (Figure 5).

**FIGURE 5.**
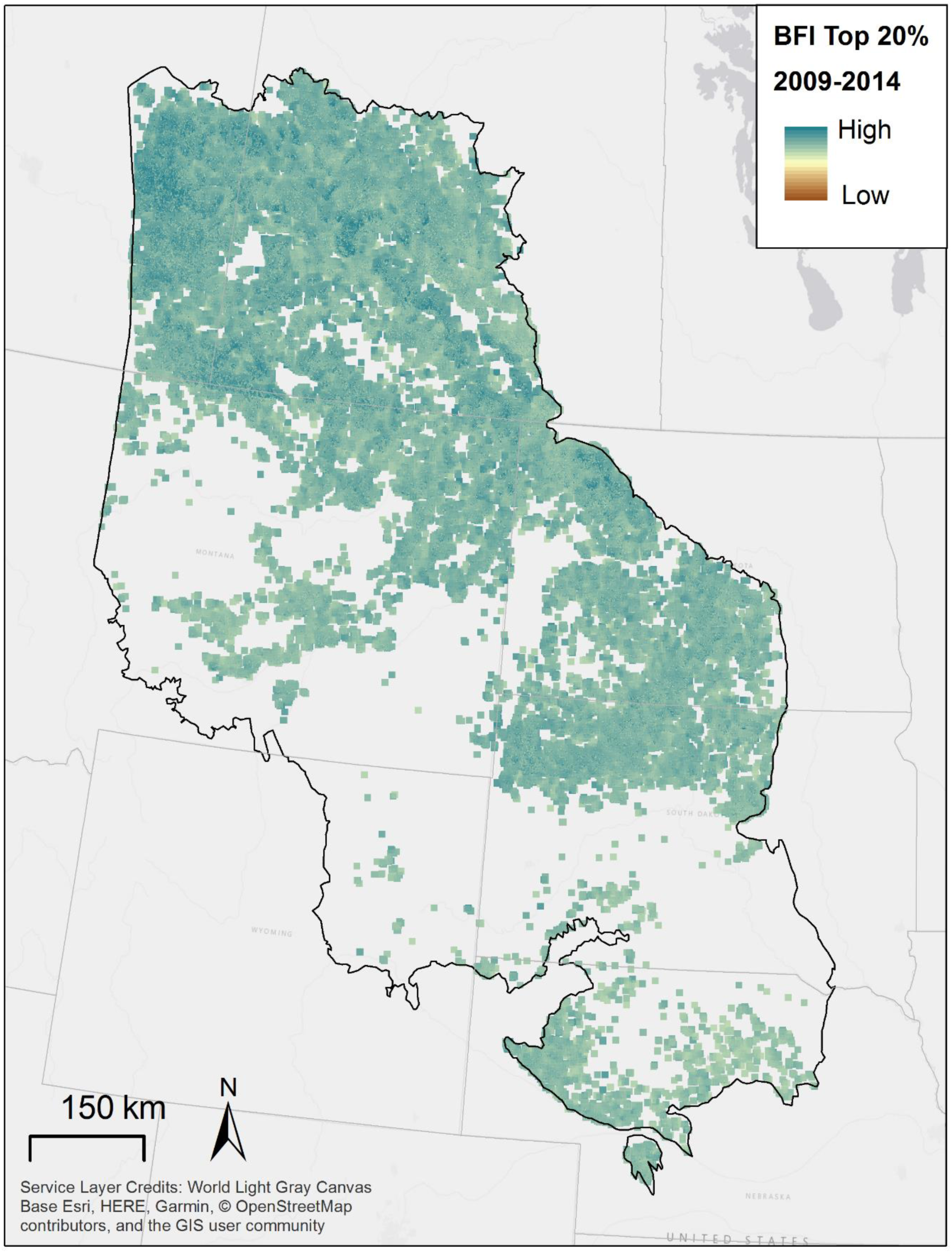
Results of spatial prioritization showing areas with BFI values in the top 20% in all years, indicating regions with the highest grassland and aridland bird community resiliency.

### 3.3 Temporal trends in relative grassland and aridland bird community resilience

BFI values were consistent over time across the Northern Great Plains study area, with mean values of 0.50 ± 2.0 e^−4^ SE in each year (slope = 8.9e^−5^ ± 5.4e^−5^, *p* = 0.00; Table 2). However, the best-fitting model had an interaction between year and strata, indicating that both mean BFI values and temporal trends varied between strata (Table 2). Montana-10, Montana-11, Montana-17, Wyoming-10, and Alberta-11 each had relatively stable BFI trends. Meanwhile, Nebraska-19 and Wyoming-18 showed slight increases, and South Dakota-17 and Wyoming-17 showed slight declines (Figure 6).

**FIGURE 6.**
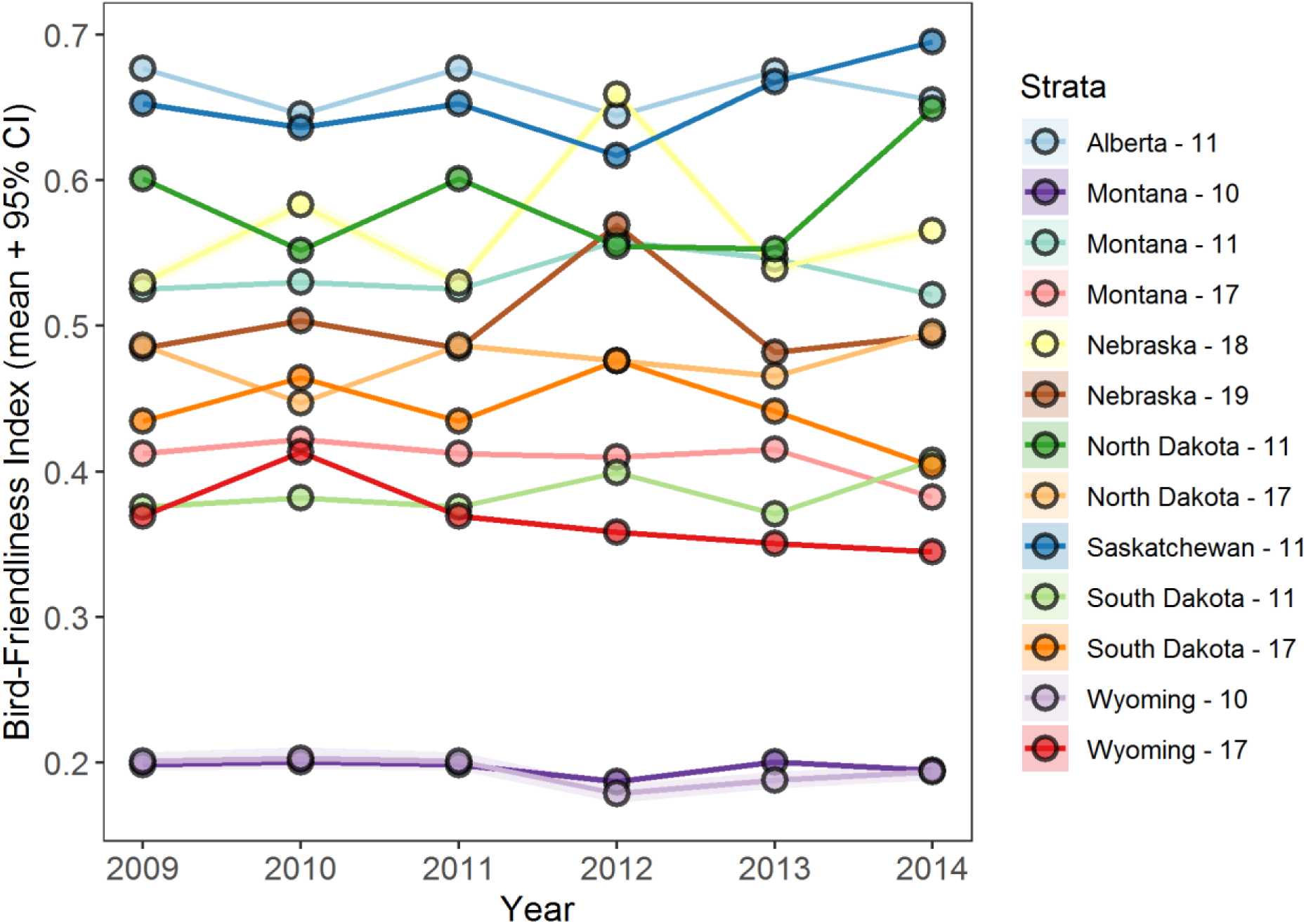
Temporal trends in BFI among strata, defined as unique combinations of states/provinces and Bird Conservation Regions. Mean (points) and 95% confidence intervals (ribbons) are shown. Locations of strata are shown in Figure 1.

### 3.4 Case study: effects of land management practices on grassland and aridland bird communities

BFI values representing grassland bird community response to simulated management were 11% higher in 2014 than BFIs estimated from observed data (Figure 7). Additionally, BFI values with simulated bird-friendly habitat management significantly increased over the six-year period (slope = 0.036 ± 0.005 SE, *p* = 0.002), while estimated BFI values (without bird-friendly habitat management) did not change during the same time period (slope = 0.014 ± 0.007 SE, *p* = 0.125). This suggests that practices recommended for use in bird-friendly grassland and aridland habitat management plans will increase the abundance and resilience of the bird community, and will be detected using the BFI.

**FIGURE 7.**
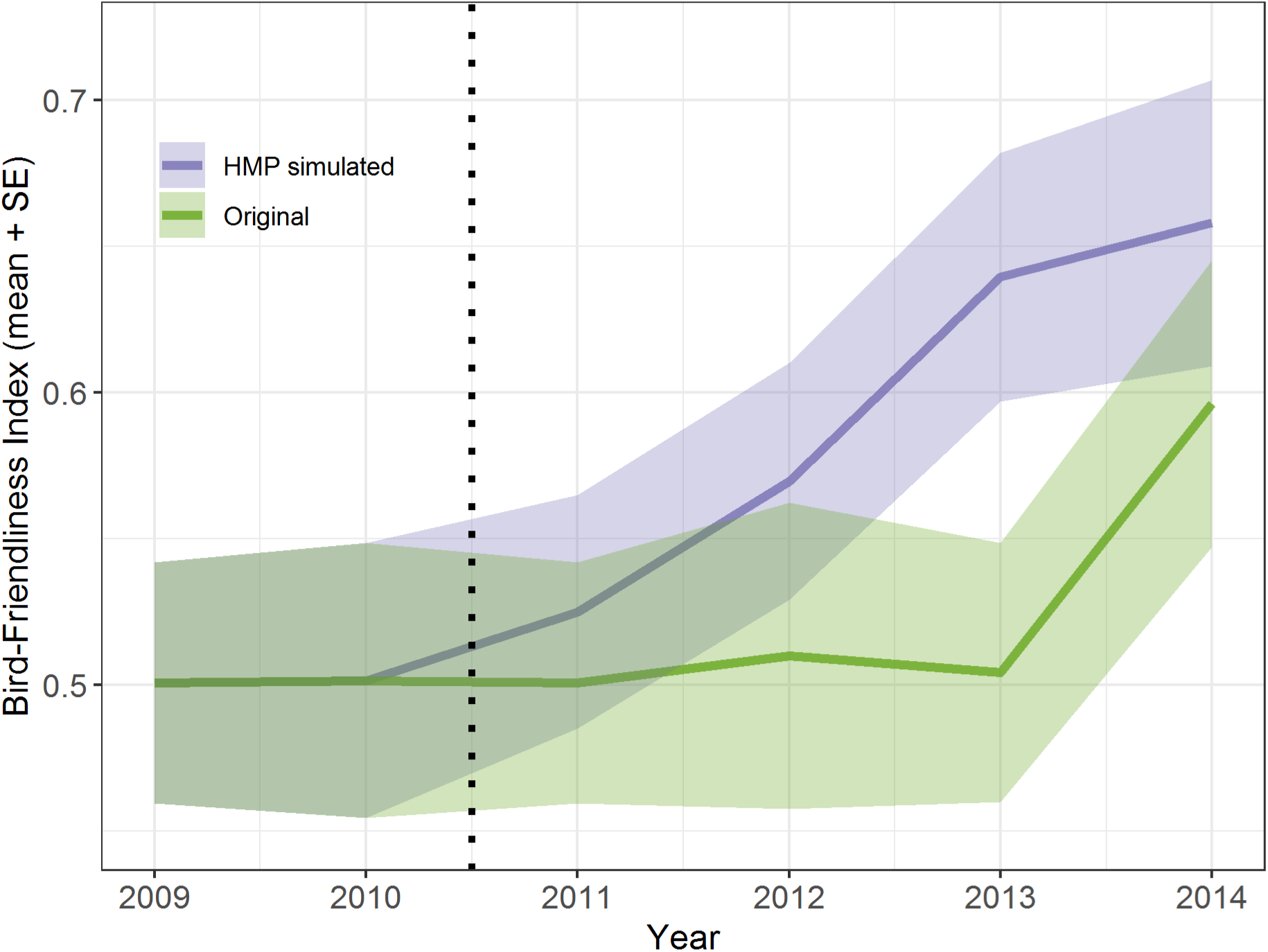
Case study showing estimated and simulated BFIs for a hypothetical management area within the northern Great Plains. Simulated BFIs representing the response to hypothetical bird-friendly habitat management implemented during 2011-2014 were higher overall than estimated BFIs from observed data, and increased over time. Mean (lines) and SE (ribbons) are shown.

## 4 DISCUSSION

The ability to quantify the impacts and success of conservation and management actions is crucial to the adaptive management cycle (Nichols & Williams, 2006). The Bird-Friendliness Index (BFI) can provide accountability and transparency for implementation of grassland and aridland habitat enhancement. The BFI evaluates the response of the entire grassland bird community to habitat management, as well as information on individual species, in an annual basis. This, in turn, can inform selection or adaptation of habitat management practices for the subsequent year, thus informing the adaptive management process.

The most resilient grassland and aridland bird communities were found in the Prairie Potholes region. The Prairie Potholes have long been known as a waterfowl nursery, but they also provide crucial habitat for grassland birds (Prairie Pothole Joint Venture, 2017). Our spatial prioritization using the BFI was consistent with other recent prioritizations using different methods, datasets, and assumptions. Prairie Pothole regions in the top 20% of BFI values coincide with Grassland Potential Conservation Areas identified in the N Great Plains Fescue Mixed-grass and NW Great Plains Mixed-grass Prairie grasslands, while the pocket of high-priority habitat in Nebraska aligns with the W Great Plains Sand Prairie Grassland Potential Conservation Area (Comer et al., 2018). Similarly, many Grassland Priority Conservation Areas (Pool & Panjabi, 2011) were identified across the Prairie Potholes, Dakotas, as well as in a few isolated pockets in northeastern Wyoming coinciding with top-priority areas identified by the BFI. A recent spatial prioritization analyzing current and future climate suitability also identified the Prairie Potholes and North Dakota as high-priority areas, similar to our findings (Grand, Wilsey, Wu, & Michel, 2019).

Conversely, the lowest BFI values were found in the southwestern regions of the study area, notably southern Montana, Wyoming, and southern South Dakota. This finding is consistent with a recent grasslands prioritization based primarily on climate suitability (Grand et al., 2019). These regions tend to be drier with more sparse vegetation, as well as lower soil moisture and productivity. While these regions provide critical habitat for aridland bird species such as Greater Sage-grouse (Burkhalter et al., 2018; Connelly, Hagen, & Schröder, 2011), they are less suitable for grassland birds that prefer lusher and more productive habitats (Fedy et al., 2018; Fisher & Davis, 2010; Harrower et al., 2017; Renfrew et al., 2013). The BFI was calculated using data for 26 predominantly grassland-breeding birds and 12 predominantly aridland-breeding birds. Thus BFI values were lower in this region overall as many grassland birds have higher mean densities than aridland birds (Table 1), and the BFI is most sensitive to changes in bird densities (Figure 4).

Evaluation of the drivers of occurrence and abundance revealed that moisture availability, terrain ruggedness, and availability of suitable habitat (grassland, shrubland) were the most important predictors. This conclusion fits well in line with what we already know of the Northern Great Plains ecosystem and grassland bird biology. The Northern Great Plains are characterized by high variability in terms of moisture, undergoing periodic bouts of large-scale drought or decreased precipitation particularly in southern and western regions, which in turn has large-scale impacts on vegetation growth (Laird, Fritz, Maasch, & Cumming, 1996). Both short-term (∼2-3 years) and long-term (24 years) studies have shown large impacts on many grassland bird species, with most responding negatively to decreases in annual precipitation and moisture levels, indicating their sensitivity to drought (George et al., 1992; Gorzo et al., 2016; Niemuth et al., 2008).

Grassland birds are also associated with low elevation sites with limited topographical variation (Wiens, 1973; Niemuth et al., 2017). In addition to being dry, the western region of the Northern Great Plains study area has greater variability in terrain and more high elevation sites, contributing to the lower BFI values. Finally, the importance of grassland, shrubland, and cropland cover in explaining variation in bird distribution and abundance is consistent with grassland and aridland bird ecology. Grassland cover is an important predictor of grassland birds associated with higher occurrence rates and densities, while shrubland and cropland cover are associated with lower occurrence rates and densities (Fedy et al., 2018; Fisher & Davis, 2010). Conversely, many aridland species such as Greater Sage-grouse, Bell’s Vireo, and Loggerhead Shrike are positively associated with shrubland habitats, increasing the importance of this predictor (Budnik, Thompson, & Ryan, 2002; Connelly et al., 2011). We were unable to incorporate annual landcover in this case study due the lack of robust annual landcover classifications that span the US / Canada border. However, future applications of the BFI should take advantage of annual landcover data, where available, because of these inherent sensitivities.

The BFI was designed for spatial comparisons within a single time period, or evaluating site-level trends in resiliency relative to other sites or the region as a whole, but not for evaluating population trends at the species or region level. As such, the standardization procedure was implemented, which resulted in relatively stable BFI values over time across strata or the study area as a whole, despite interannual variation in grassland and aridland bird density. Standardization enables isolation of local-scale effects, such as bird response to habitat management or regional differences in land use, from large-scale processes such as long-term population-level declines due to shared drivers such as climate oscillations or wintering ground effects (Gorzo et al., 2016; Macías-Duarte et al., 2017). The effects of this standardization procedure were apparent in the habitat management case study. Though neither mean BFI across the study area nor mean observed BFI at the case study site increased over six years, simulated habitat management increased grassland and aridland bird densities and diversity relative to the surrounding landscape, producing a significant increasing trend in BFI. However, the standardization step could be removed for studies with an objective of tracking long-term trends in ecological resilience across large spatial scales, rather than evaluating local response to conservation actions.

Conservation efforts should aim to do more than prevent the extinction of species, but rather should be aimed at preventing species from becoming threatened in the first place, as well as provide conditions that enhance ecosystem processes (Rodrigues, 2006). By identifying early on which species and communities are doing well or poorly, and where, the BFI can pinpoint priority strongholds for conservation, or opportunities for restoration, both of which may contribute to population stabilization or even growth. To be able to do this effectively requires indicators that are rigorous, repeatable, and easily understood (Balmford et al., 2005). National Audubon Society deploys the BFI as an accountability metric for the Conservation Ranching Program (https://www.audubon.org/conservation/ranching), a market-based conservation solution that aims to conserve imperiled grassland and aridland bird species and habitats through partnership with private landowners. Other public/private grassland and aridland conservation efforts, e.g., World Wildlife Fund’s Grasslands program (https://www.worldwildlife.org/habitats/grasslands), Sodsaver (enacted as part of the Agricultural Act of 2014) and Natural Resources Conservation Service’s EQIP (https://www.nrcs.usda.gov/wps/portal/nrcs/main/national/programs/financial/eqip/) may also benefit from the accountability provided by this community ecological resilience metric. The BFI is a tool by which conservationists and managers can carry out accountable conservation now and into the future.

## DATA ACCESSIBILITY STATEMENT

The R scripts used to conduct the simulations and analyses will be made available through Data Dryad upon manuscript acceptance.

