## Supporting Tables and Figures for "Metrics for conservation success: using the ‘Bird-Friendliness Index’ to evaluate grassland and aridland bird community resilience across the Northern Great Plains ecosystem"

**TABLE S1.** Variables used as predictors in the species distribution models and their sources and citations.

| Type | Variable | Source |
|---|---|---|
| Climate | Climatic moisture deficit | Climatic Research Unit Timeseries 3.22 dataset, 1981-2010 climate normal (Wang et al., 2016) |
| Climate | Spring degree-days below 0°C | Climatic Research Unit Timeseries 3.22 dataset, 1981-2010 climate normal (Wang et al., 2016) |
| Ecosystem | Litter biomass (g $C/m^2$) | CASA ecosystem model (Potter et al. 1993, 2007) |
| Ecosystem | Net primary productivity (g $C/m^2$) | CASA ecosystem model (Potter et al. 1993, 2007) |
| Ecosystem | Nitrous oxide flux (g $N_2O/m^2$-day) | CASA ecosystem model (Potter et al. 1993, 2007) |
| Ecosystem | Soil moisture (cm; 0-10 cm depth) | CASA ecosystem model (Potter et al. 1993, 2007) |
| Ecosystem | Surface air temperature (°C) | CASA ecosystem model (Potter et al. 1993, 2007) |
| Ecosystem | Total $CO_2$ (g) | CASA ecosystem model (Potter et al. 1993, 2007) |
| Ecosystem | Evaporation | CASA ecosystem model (Potter et al. 1993, 2007) |
| Landcover | Proportion cropland | Derived from Commission for Environmental Cooperation's North American Environmental Atlas 2010 (Canada Centre for Remote Sensing (CCRS) et al. 2013) |
| Landcover | Proportion grassland | Derived from Commission for Environmental Cooperation's North American Environmental Atlas 2010 (Canada Centre for Remote Sensing (CCRS) et al. 2013) |

| | | |
|---|---|---|
| Landcover | Proportion shrubland | Derived from Commission for Environmental Cooperation’s North American Environmental Atlas 2010 (Canada Centre for Remote Sensing (CCRS) et al. 2013) |
| Landcover | Proportion wetland | Derived from Commission for Environmental Cooperation’s North American Environmental Atlas 2010 (Canada Centre for Remote Sensing (CCRS) et al. 2013) |
| Landcover | Grassland mean patch area | Derived from Commission for Environmental Cooperation’s North American Environmental Atlas 2010 (Canada Centre for Remote Sensing (CCRS) et al. 2013) |
| Landcover | Grassland patch cohesion index | Derived from Commission for Environmental Cooperation’s North American Environmental Atlas 2010 (Canada Centre for Remote Sensing (CCRS) et al. 2013) |
| Temporal | Year | Year |
| Topographical | Terrain ruggedness index | Derived from digital elevation model (Riley et al. 1999) |

**TABLE S2.** Model fit statistics for presence/absence and abundance models for 34 species across the Northern Great Plains during 2009-2014. Statistics include AUC (presence/absence only), TSS (presence/absence only), cross-validated deviance explained (Dev. Exp.), cross-validated correlation (Corr.), and the Moran's I test for residual spatial autocorrelation. Model fit statistics were calculated with 12-fold cross-validation, and are averaged across 25 bootstrapped geographically filtered datasets (mean ± SE reported).

| | Presence/absence model | | | | | Abundance model | | |
|---|---|---|---|---|---|---|---|---|
| Common Name | AUC | TSS | Dev. Exp. | Corr. | Moran's I | Dev. Exp. | Corr. | Moran's I |
| Baird's Sparrow | 0.74 ± 0.05 | 0.18 ± 0.01 | 0.42 ± 0.01 | 0.26 ± 0.04 | 0.00 ± 0.00 | 0.60 ± 0.02 | 0.20 ± 0.06 | 0.00 ± 0.00 |
| Bell's Vireo | 0.53 ± 0.08 | 0.05 ± 0.00 | 0.41 ± 0.00 | 0.11 ± 0.06 | 0.04 ± 0.00 | 0.01 ± 0.00 | 0.12 ± 0.02 | -0.01 ± 0.00 |
| Bobolink | 0.74 ± 0.04 | 0.17 ± 0.00 | 0.42 ± 0.00 | 0.32 ± 0.08 | 0.00 ± 0.00 | 0.52 ± 0.01 | 0.32 ± 0.09 | 0.01 ± 0.00 |
| Brewer's Sparrow | 0.74 ± 0.05 | 0.32 ± 0.02 | 0.55 ± 0.01 | 0.30 ± 0.07 | 0.00 ± 0.00 | 0.50 ± 0.01 | 0.08 ± 0.08 | 0.01 ± 0.00 |
| Burrowing Owl | 0.87 ± 0.04 | 0.11 ± 0.00 | 0.42 ± 0.00 | 0.36 ± 0.08 | 0.05 ± 0.00 | 0.00 ± 0.00 | 0.36 ± 0.09 | 0.04 ± 0.00 |
| Canyon Wren | 0.79 ± 0.04 | 0.03 ± 0.00 | 0.34 ± 0.00 | 0.16 ± 0.05 | 0.05 ± 0.00 | 0.00 ± 0.00 | 0.16 ± 0.04 | 0.05 ± 0.00 |
| Chestnut-collared Longspur | 0.76 ± 0.03 | 0.27 ± 0.01 | 0.47 ± 0.01 | 0.33 ± 0.05 | 0.00 ± 0.00 | 0.58 ± 0.01 | 0.29 ± 0.06 | 0.00 ± 0.00 |
| Clay-colored Sparrow | 0.64 ± 0.06 | 0.24 ± 0.01 | 0.25 ± 0.00 | 0.16 ± 0.07 | 0.00 ± 0.00 | 0.62 ± 0.00 | 0.17 ± 0.09 | 0.00 ± 0.00 |
| Dickcissel | 0.67 ± 0.07 | 0.12 ± 0.00 | 0.35 ± 0.01 | 0.18 ± 0.06 | 0.00 ± 0.00 | 0.35 ± 0.05 | 0.09 ± 0.04 | 0.00 ± 0.00 |
| Eastern Kingbird | 0.70 ± 0.04 | 0.45 ± 0.00 | 0.26 ± 0.00 | 0.33 ± 0.07 | 0.00 ± 0.00 | 0.27 ± 0.01 | 0.28 ± 0.07 | 0.00 ± 0.00 |
| Ferruginous Hawk | 0.61 ± 0.08 | 0.02 ± 0.00 | 0.08 ± 0.02 | 0.09 ± 0.07 | 0.00 ± 0.00 | 0.08 ± 0.01 | 0.15 ± 0.06 | 0.00 ± 0.00 |
| Grasshopper Sparrow | 0.82 ± 0.05 | 0.60 ± 0.01 | 0.56 ± 0.00 | 0.53 ± 0.09 | 0.00 ± 0.00 | 0.67 ± 0.00 | 0.35 ± 0.10 | 0.00 ± 0.00 |
| Greater Prairie-chicken | 0.86 ± 0.06 | 0.17 ± 0.02 | 0.83 ± 0.01 | 0.50 ± 0.12 | 0.05 ± 0.00 | 0.16 ± 0.00 | 0.21 ± 0.12 | 0.02 ± 0.00 |
| Greater Sage-grouse | 0.71 ± 0.04 | 0.06 ± 0.01 | 0.39 ± 0.02 | 0.13 ± 0.04 | 0.00 ± 0.00 | 0.04 ± 0.00 | 0.02 ± 0.03 | 0.00 ± 0.00 |
| Green-tailed Towhee | 0.59 ± 0.02 | 0.05 ± 0.00 | 0.41 ± 0.01 | 0.15 ± 0.07 | 0.00 ± 0.00 | 0.22 ± 0.05 | 0.23 ± 0.07 | 0.00 ± 0.00 |
| Horned Lark | 0.82 ± 0.05 | 0.54 ± 0.01 | 0.49 ± 0.01 | 0.54 ± 0.09 | 0.00 ± 0.00 | 0.48 ± 0.00 | 0.48 ± 0.07 | 0.01 ± 0.00 |
| Lark Bunting | 0.89 ± 0.00 | 0.47 ± 0.01 | 0.38 ± 0.00 | 0.54 ± 0.07 | 0.00 ± 0.00 | 0.46 ± 0.01 | 0.36 ± 0.07 | 0.00 ± 0.00 |
| Lark Sparrow | 0.71 ± 0.05 | 0.47 ± 0.01 | 0.37 ± 0.00 | 0.35 ± 0.08 | 0.00 ± 0.00 | 0.41 ± 0.01 | 0.34 ± 0.04 | 0.00 ± 0.00 |
| Loggerhead Shrike | 0.62 ± 0.06 | 0.12 ± 0.00 | 0.19 ± 0.01 | 0.12 ± 0.06 | 0.00 ± 0.00 | 0.04 ± 0.02 | 0.10 ± 0.05 | 0.00 ± 0.00 |
| Long-billed Curlew | 0.83 ± 0.02 | 0.15 ± 0.01 | 0.45 ± 0.01 | 0.43 ± 0.05 | 0.00 ± 0.00 | 0.36 ± 0.00 | 0.19 ± 0.06 | 0.00 ± 0.00 |
| McCown's Longspur | 0.56 ± 0.06 | 0.04 ± 0.00 | 0.35 ± 0.01 | 0.00 ± 0.02 | 0.00 ± 0.00 | 0.01 ± 0.00 | 0.02 ± 0.03 | 0.00 ± 0.00 |
| Mountain Plover | 0.97 ± 0.03 | 0.09 ± 0.00 | 0.96 ± 0.00 | 0.88 ± 0.04 | 0.04 ± 0.00 | 0.02 ± 0.00 | -0.10 ± 0.05 | 0.01 ± 0.00 |
| Northern Harrier | 0.71 ± 0.05 | 0.20 ± 0.00 | 0.22 ± 0.00 | 0.24 ± 0.05 | 0.00 ± 0.00 | 0.30 ± 0.01 | 0.17 ± 0.06 | 0.00 ± 0.00 |
| Rock Wren | 0.76 ± 0.04 | 0.29 ± 0.00 | 0.35 ± 0.01 | 0.41 ± 0.07 | 0.00 ± 0.00 | 0.06 ± 0.03 | 0.20 ± 0.07 | 0.00 ± 0.00 |

| | | | | | | | | |
|---|---|---|---|---|---|---|---|---|
| Sage Thrasher | 0.57 ± 0.07 | 0.06 ± 0.00 | 0.31 ± 0.01 | 0.00 ± 0.07 | 0.00 ± 0.00 | 0.00 ± 0.00 | 0.05 ± 0.06 | 0.00 ± 0.00 |
| Savannah Sparrow | 0.72 ± 0.05 | 0.17 ± 0.00 | 0.30 ± 0.00 | 0.28 ± 0.08 | 0.00 ± 0.00 | 0.43 ± 0.00 | 0.27 ± 0.10 | 0.00 ± 0.00 |
| Sharp-tailed Grouse | 0.65 ± 0.04 | 0.21 ± 0.01 | 0.26 ± 0.01 | 0.18 ± 0.06 | 0.00 ± 0.00 | 0.24 ± 0.03 | 0.22 ± 0.06 | 0.00 ± 0.00 |
| Sprague’s Pipit | 0.76 ± 0.03 | 0.16 ± 0.00 | 0.46 ± 0.01 | 0.34 ± 0.07 | 0.00 ± 0.00 | 0.38 ± 0.01 | 0.34 ± 0.05 | 0.00 ± 0.00 |
| Swainson's Hawk | 0.68 ± 0.06 | 0.12 ± 0.00 | 0.21 ± 0.02 | 0.15 ± 0.05 | 0.00 ± 0.00 | 0.07 ± 0.01 | 0.15 ± 0.04 | 0.00 ± 0.00 |
| Upland Sandpiper | 0.76 ± 0.05 | 0.52 ± 0.01 | 0.43 ± 0.01 | 0.45 ± 0.09 | 0.00 ± 0.00 | 0.69 ± 0.01 | 0.28 ± 0.07 | 0.00 ± 0.00 |
| Vesper Sparrow | 0.93 ± 0.00 | 0.44 ± 0.01 | 0.49 ± 0.01 | 0.31 ± 0.09 | 0.00 ± 0.00 | 0.32 ± 0.00 | 0.22 ± 0.07 | 0.00 ± 0.00 |
| Western Kingbird | 0.63 ± 0.05 | 0.36 ± 0.00 | 0.15 ± 0.00 | 0.18 ± 0.07 | 0.00 ± 0.00 | 0.51 ± 0.01 | 0.40 ± 0.06 | 0.00 ± 0.00 |
| Western Meadowlark | 0.99 ± 0.00 | 0.72 ± 0.01 | 0.79 ± 0.00 | 0.60 ± 0.11 | 0.01 ± 0.00 | 0.75 ± 0.00 | 0.58 ± 0.09 | 0.01 ± 0.00 |
| White-throated Swift | 0.68 ± 0.05 | 0.04 ± 0.00 | 0.14 ± 0.01 | 0.15 ± 0.06 | 0.00 ± 0.00 | 0.26 ± 0.02 | 0.12 ± 0.08 | 0.00 ± 0.00 |

### FIGURES

**FIGURE S1.** Mean variable importance and 95% confidence intervals for 17 variables used as predictors in presence/absence (A) and abundance (B) species distribution models for Baird's Sparrow across the Northern Great Plains during 2009-2014.

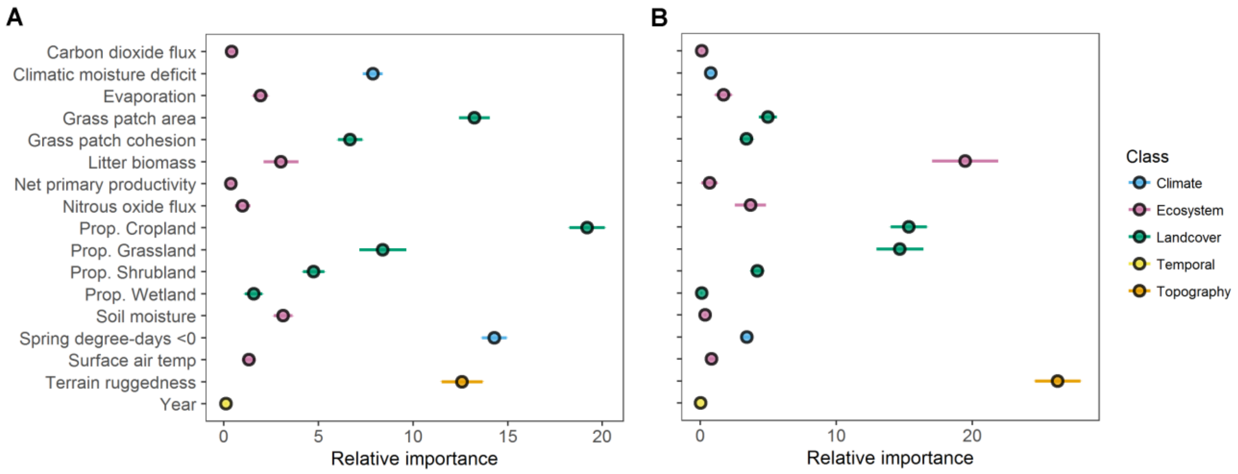


**FIGURE S2.** Mean variable importance and 95% confidence intervals for 17 variables used as predictors in presence/absence (A) and abundance (B) species distribution models for Bell's Vireo across the Northern Great Plains during 2009-2014.

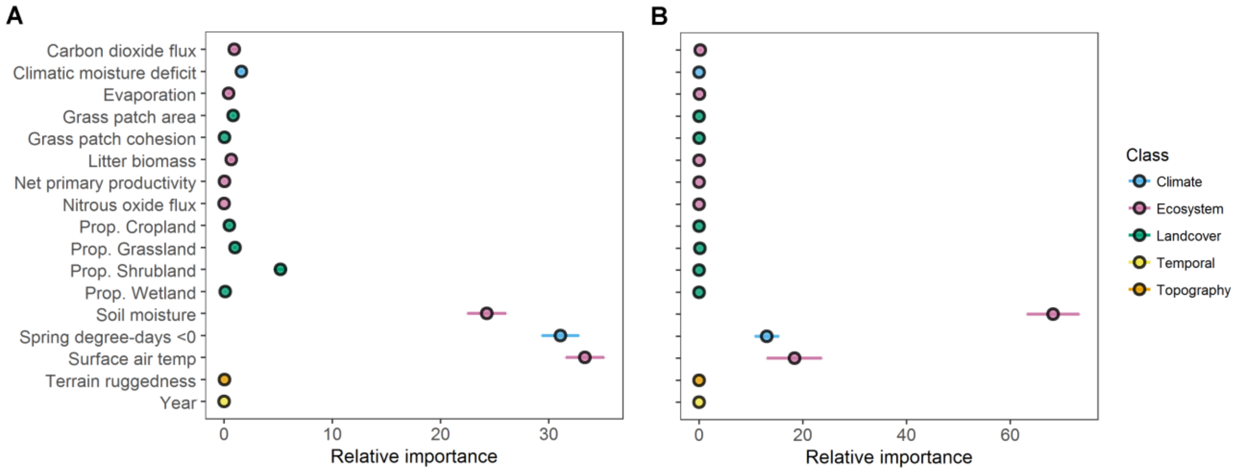

**FIGURE S3.** Mean variable importance and 95% confidence intervals for 17 variables used as predictors in presence/absence (A) and abundance (B) species distribution models for Bobolink across the Northern Great Plains during 2009-2014.

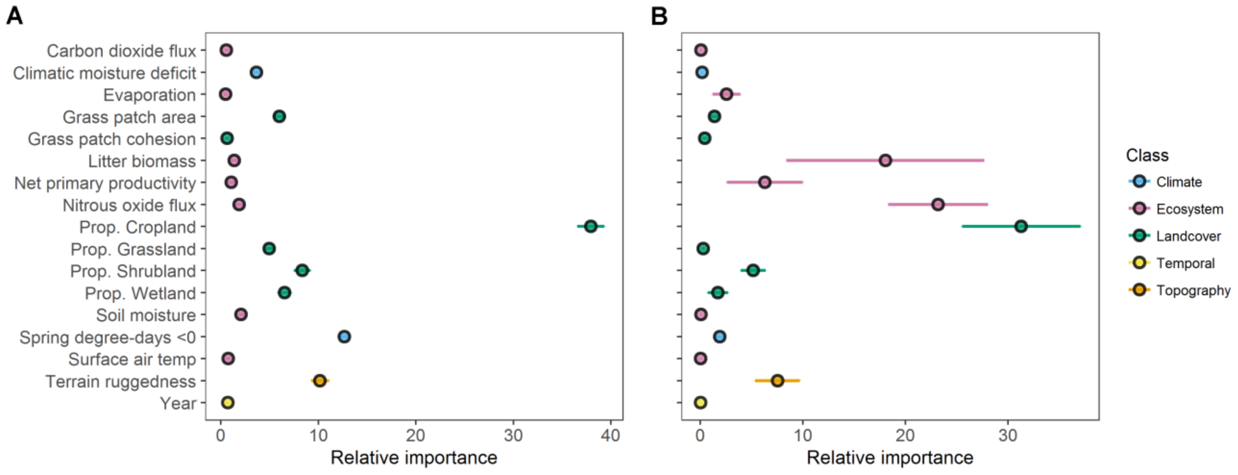


**FIGURE S4.** Mean variable importance and 95% confidence intervals for 17 variables used as predictors in presence/absence (A) and abundance (B) species distribution models for Brewer's Sparrow across the Northern Great Plains during 2009-2014.

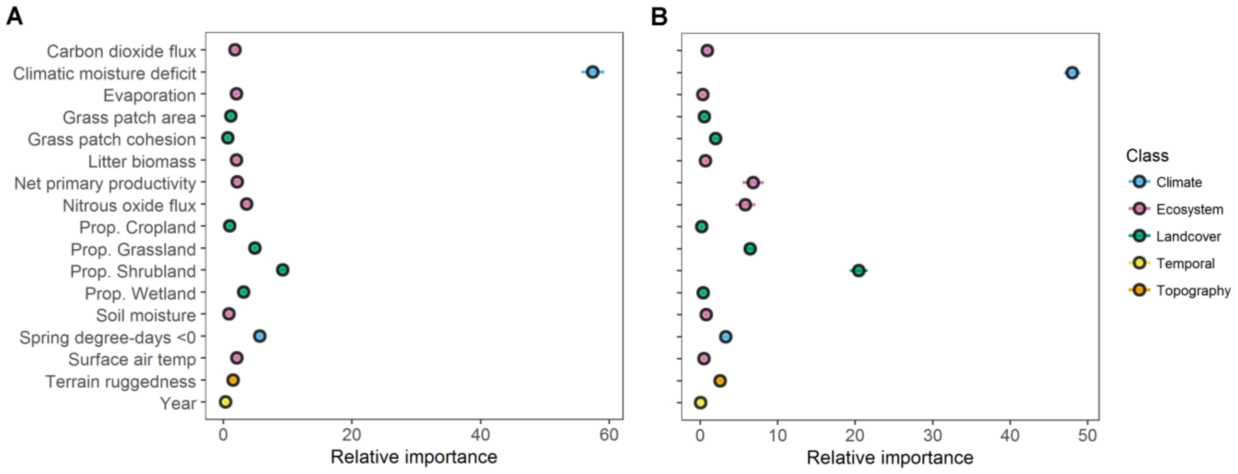

**FIGURE S5.** Mean variable importance and 95% confidence intervals for 17 variables used as predictors in presence/absence (A) and abundance (B) species distribution models for Burrowing Owl across the Northern Great Plains during 2009-2014.

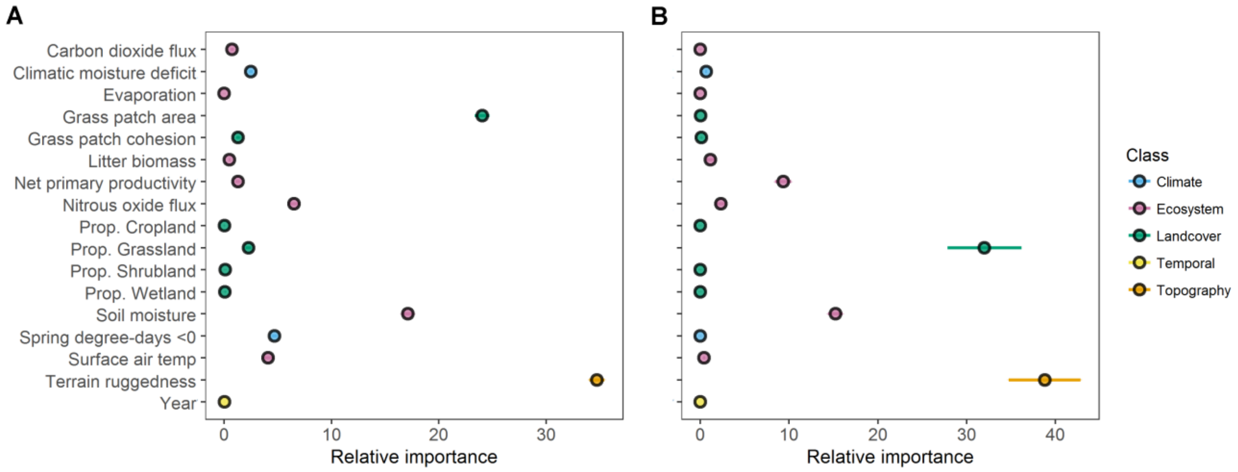


**FIGURE S6.** Mean variable importance and 95% confidence intervals for 17 variables used as predictors in presence/absence (A) and abundance (B) species distribution models for Canyon Wren across the Northern Great Plains during 2009-2014.

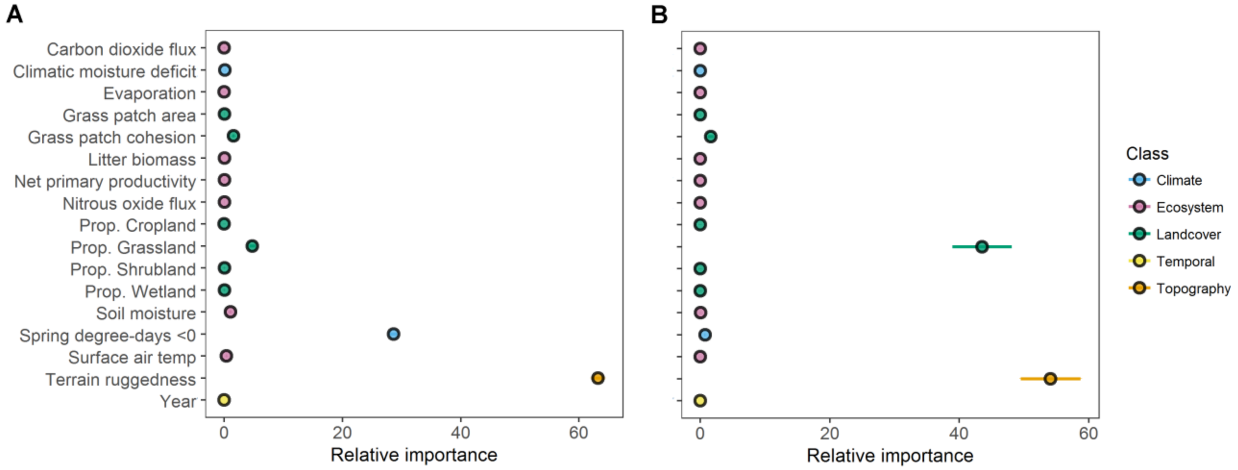

**FIGURE S7.** Mean variable importance and 95% confidence intervals for 17 variables used as predictors in presence/absence (A) and abundance (B) species distribution models for Chestnut-collared Longspur across the Northern Great Plains during 2009-2014.

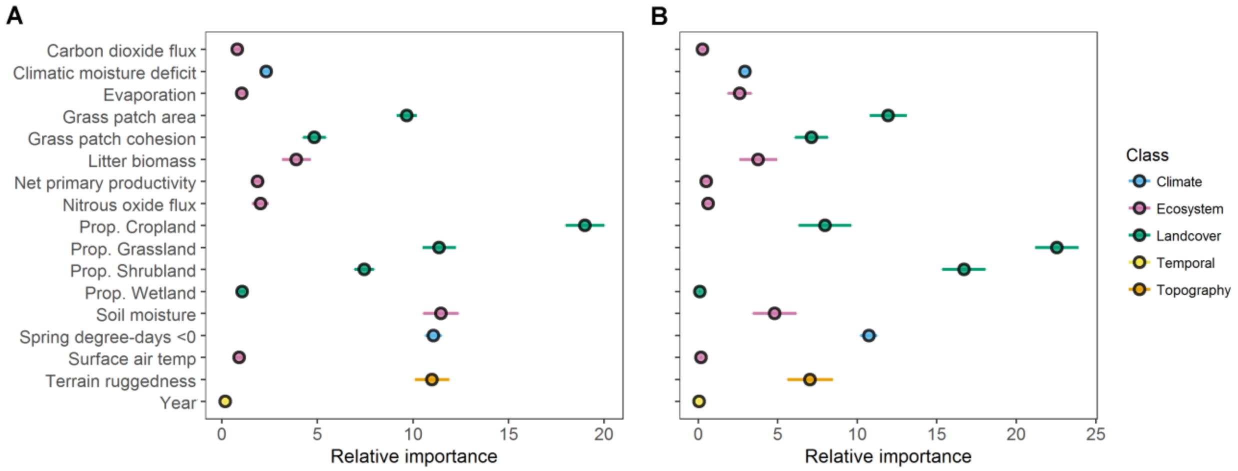


**FIGURE S8.** Mean variable importance and 95% confidence intervals for 17 variables used as predictors in presence/absence (A) and abundance (B) species distribution models for Clay-colored Sparrow across the Northern Great Plains during 2009-2014.

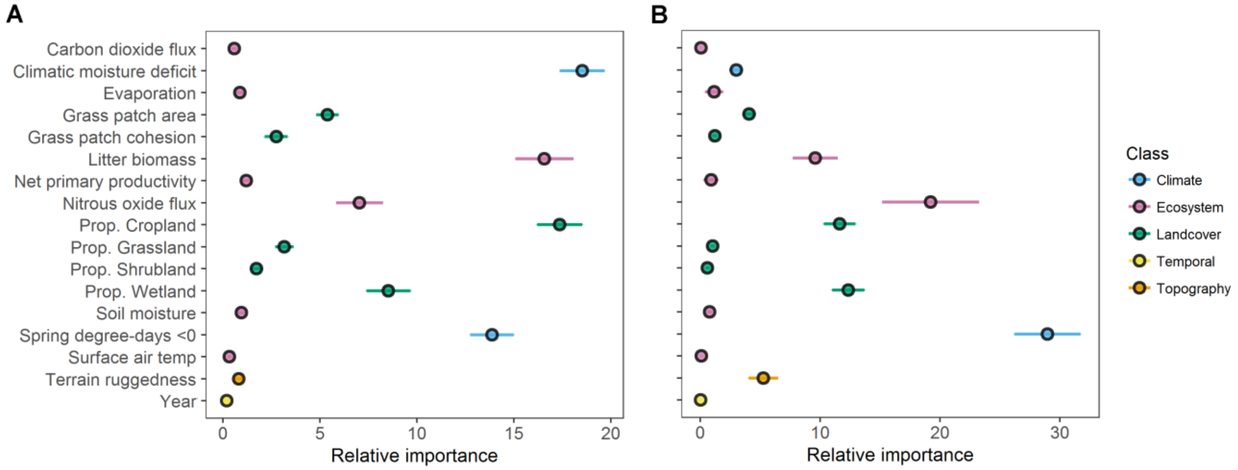

**FIGURE S9.** Mean variable importance and 95% confidence intervals for 17 variables used as predictors in presence/absence (A) and abundance (B) species distribution models for Dickcissel across the Northern Great Plains during 2009-2014.

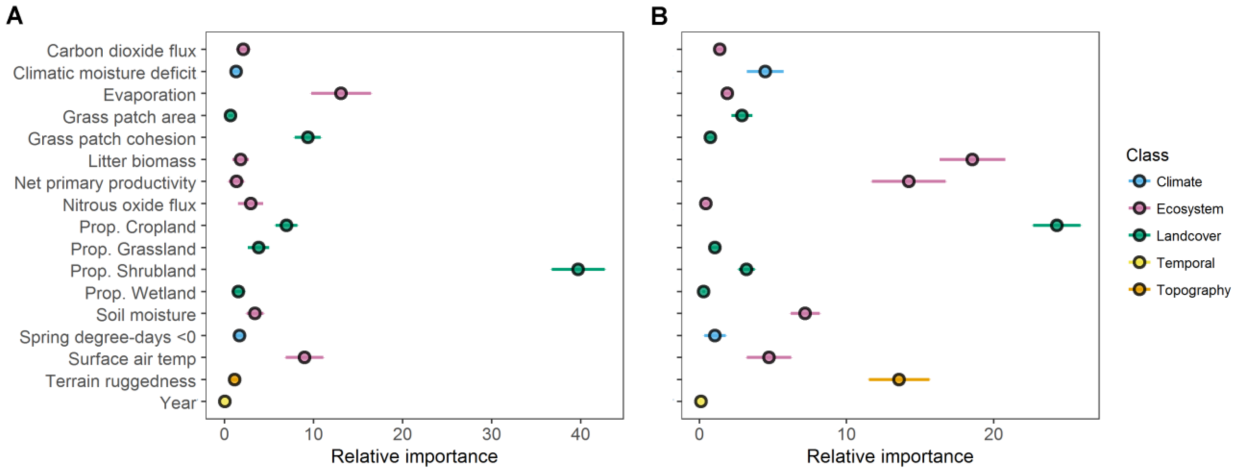


**FIGURE S10.** Mean variable importance and 95% confidence intervals for 17 variables used as predictors in presence/absence (A) and abundance (B) species distribution models for Eastern Kingbird across the Northern Great Plains during 2009-2014.

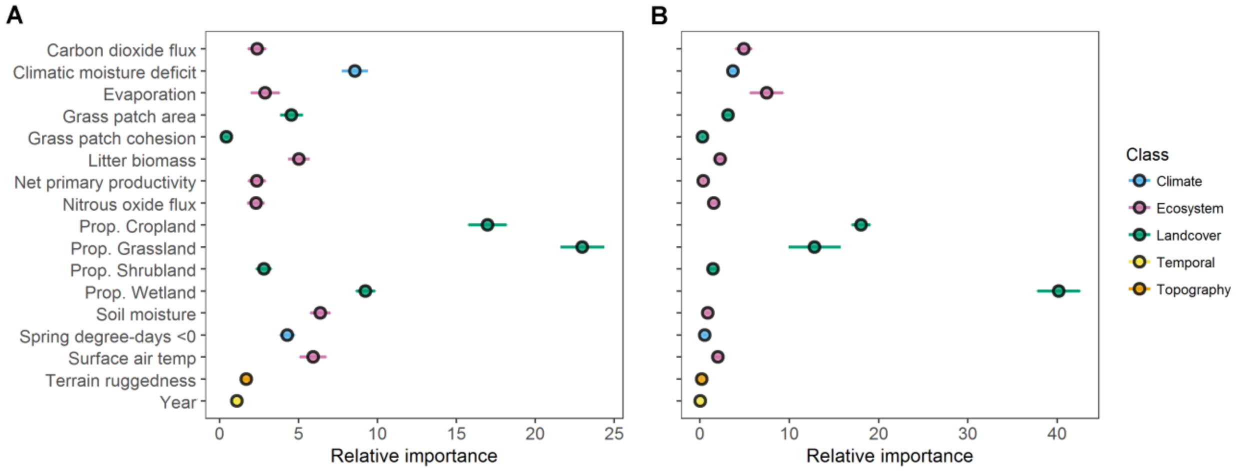

**FIGURE S11.** Mean variable importance and 95% confidence intervals for 17 variables used as predictors in presence/absence (A) and abundance (B) species distribution models for Ferruginous Hawk across the Northern Great Plains during 2009-2014.

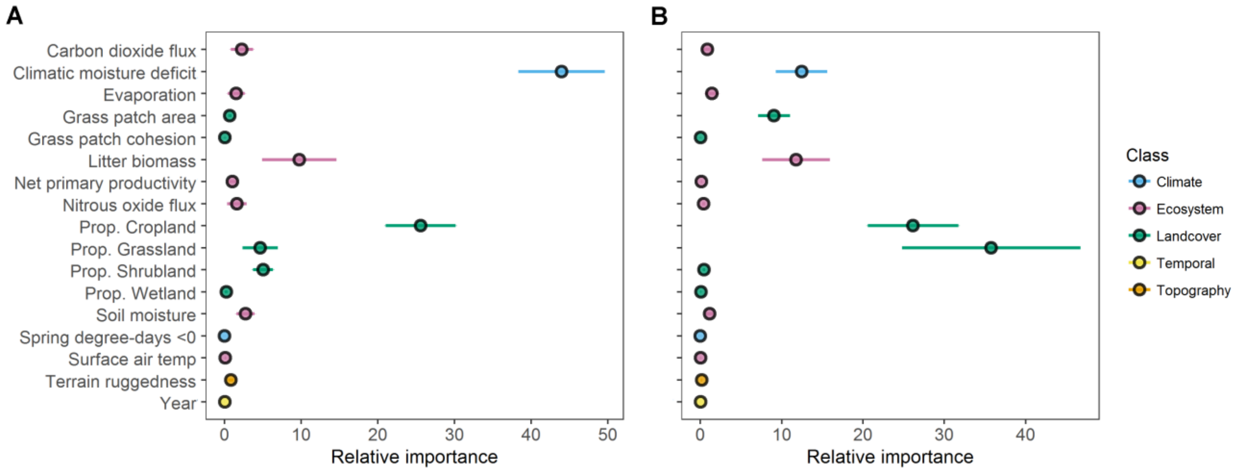


**FIGURE S12.** Mean variable importance and 95% confidence intervals for 17 variables used as predictors in presence/absence (A) and abundance (B) species distribution models for Grasshopper Sparrow across the Northern Great Plains during 2009-2014.

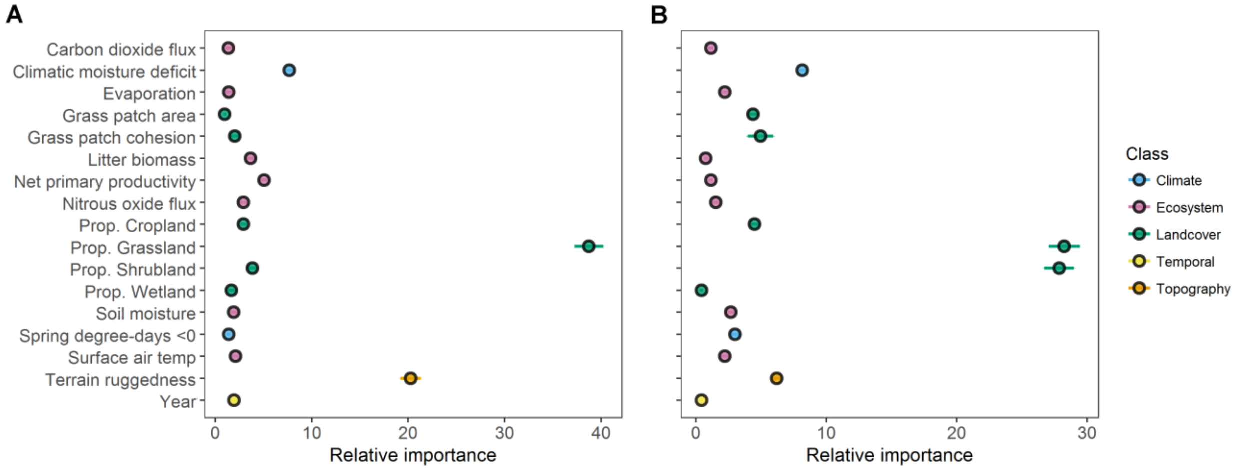

**Figure S13.** Mean variable importance and 95% confidence intervals for 17 variables used as predictors in presence/absence (A) and abundance (B) species distribution models for Greater Prairie-chicken across the Northern Great Plains during 2009-2014.

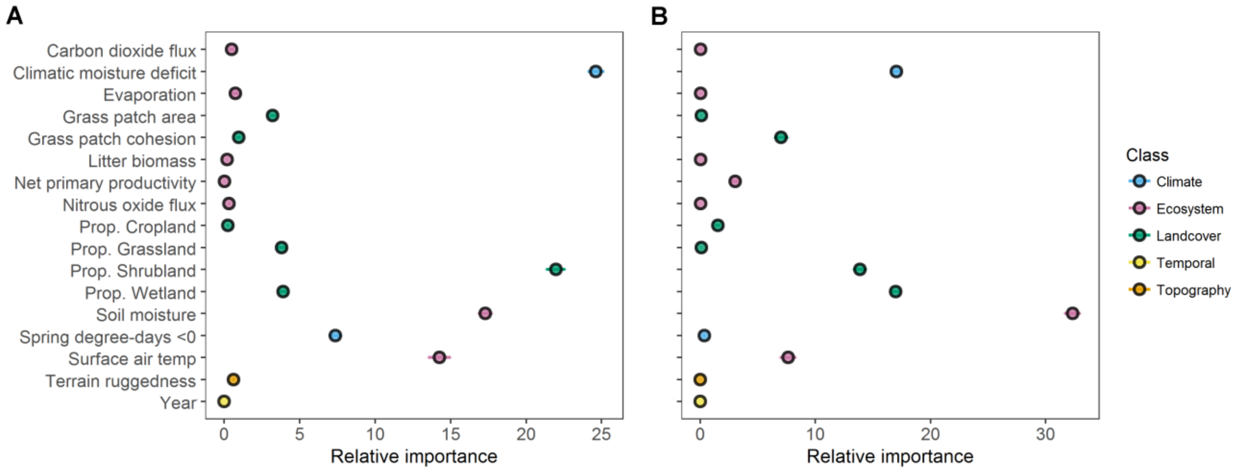


**FIGURE S14.** Mean variable importance and 95% confidence intervals for 17 variables used as predictors in presence/absence (A) and abundance (B) species distribution models for Greater Sage-grouse across the Northern Great Plains during 2009-2014.

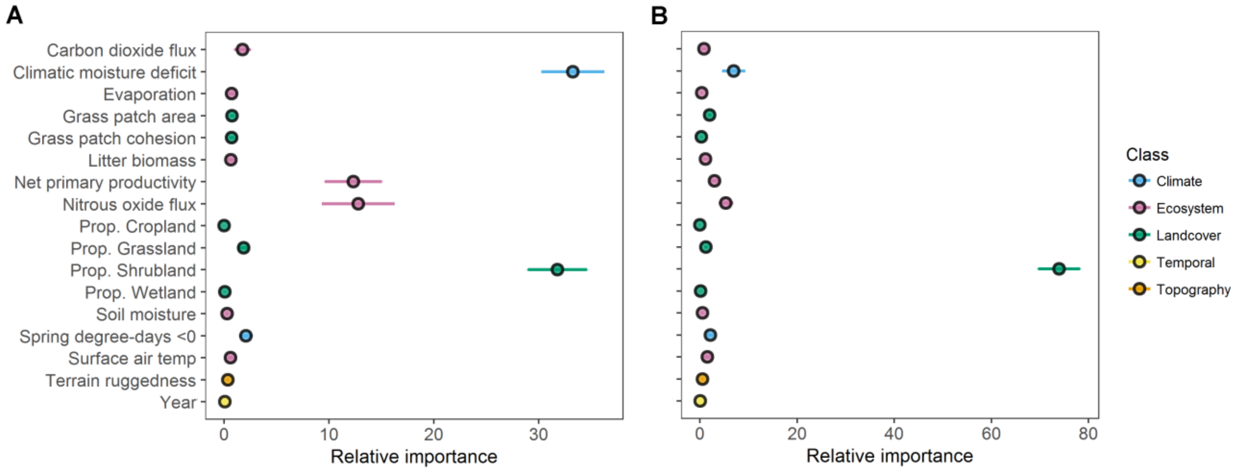

**FIGURE S15.** Mean variable importance and 95% confidence intervals for 17 variables used as predictors in presence/absence (A) and abundance (B) species distribution models for Green-tailed Towhee across the Northern Great Plains during 2009-2014.

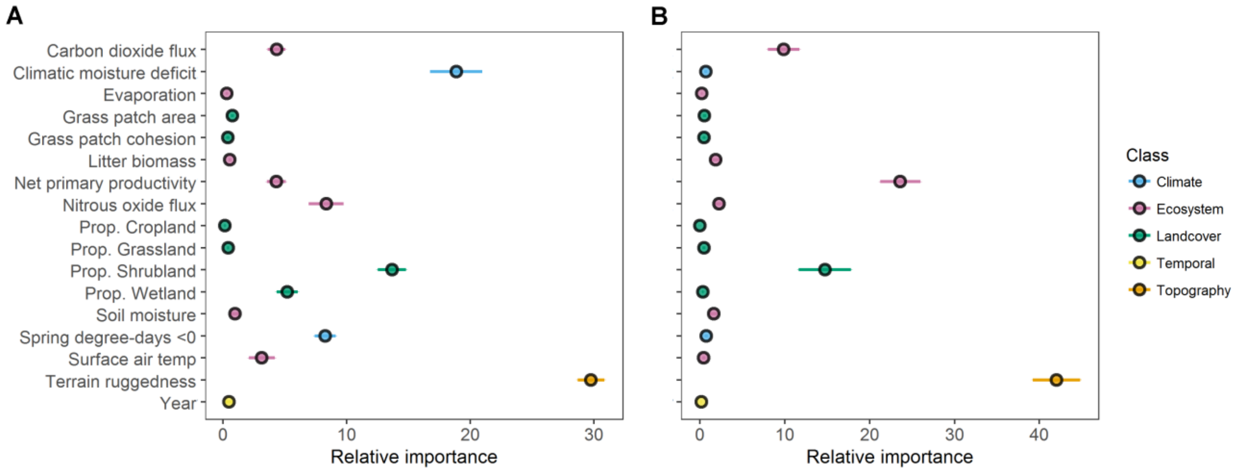


**FIGURE S16.** Mean variable importance and 95% confidence intervals for 17 variables used as predictors in presence/absence (A) and abundance (B) species distribution models for Horned Lark across the Northern Great Plains during 2009-2014.

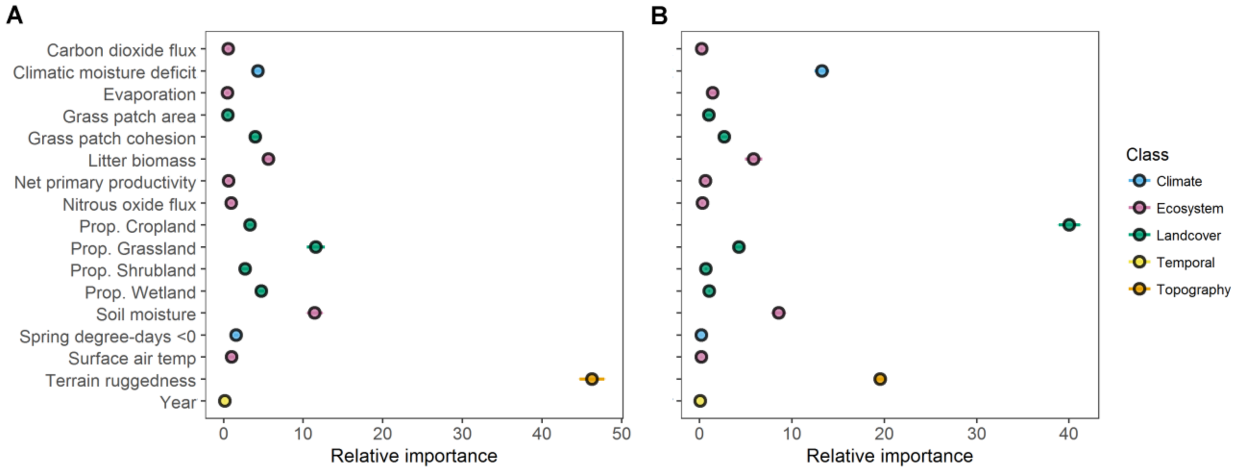

**FIGURE S17.** Mean variable importance and 95% confidence intervals for 17 variables used as predictors in presence/absence (A) and abundance (B) species distribution models for Lark Bunting across the Northern Great Plains during 2009-2014.

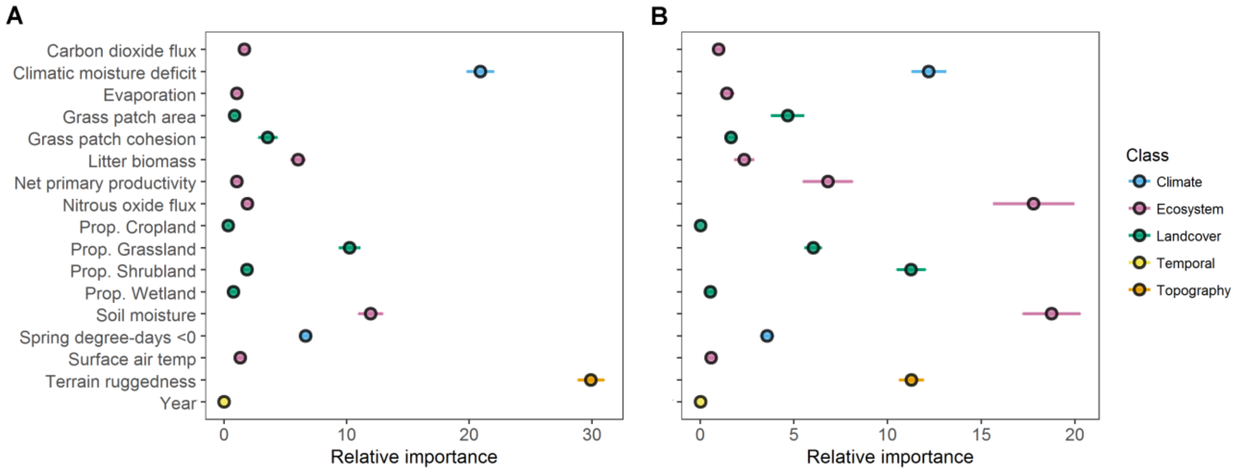


**FIGURE S18.** Mean variable importance and 95% confidence intervals for 17 variables used as predictors in presence/absence (A) and abundance (B) species distribution models for Lark Sparrow across the Northern Great Plains during 2009-2014.

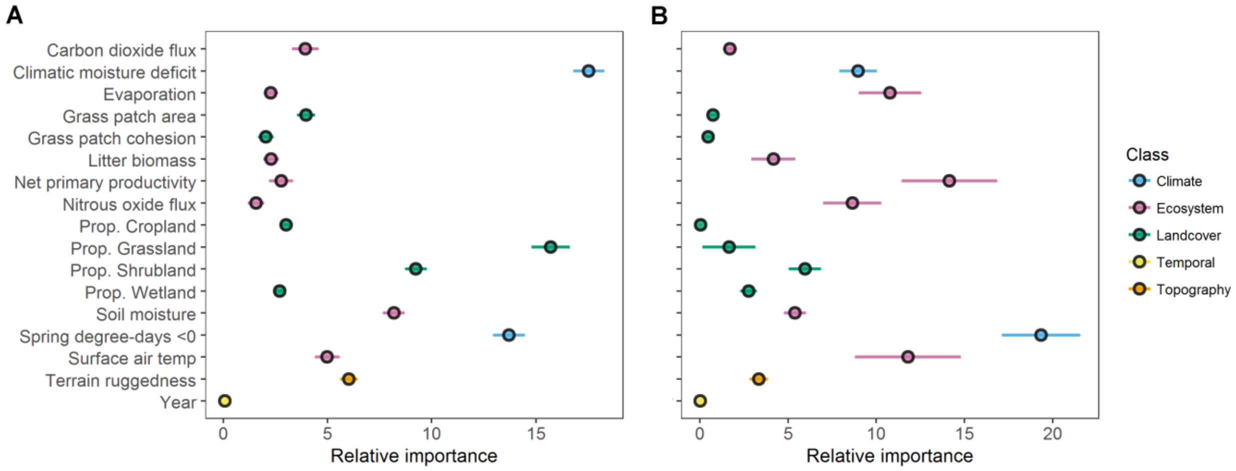

**FIGURE S19.** Mean variable importance and 95% confidence intervals for 17 variables used as predictors in presence/absence (A) and abundance (B) species distribution models for Loggerhead Shrike across the Northern Great Plains during 2009-2014.

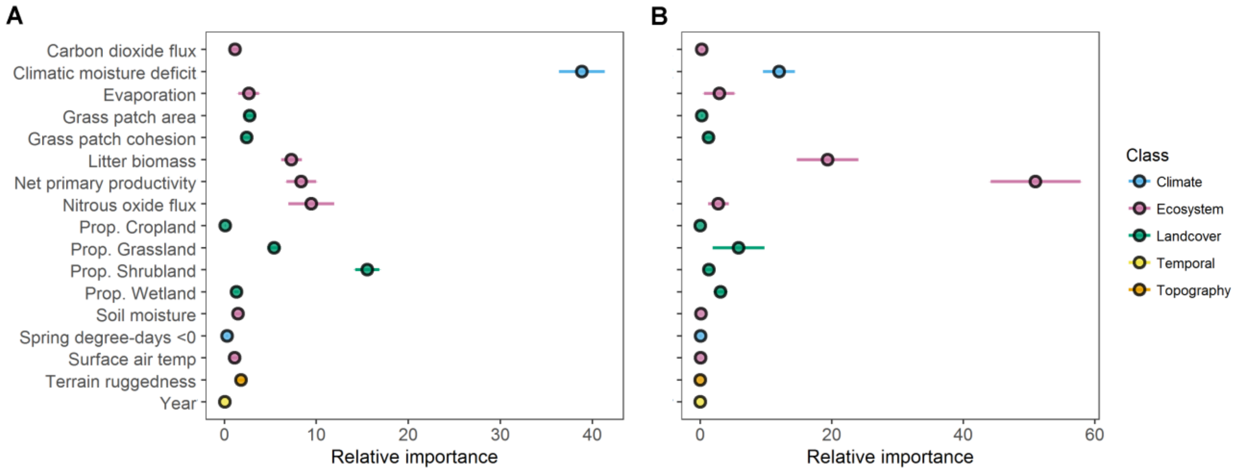


**FIGURE S20.** Mean variable importance and 95% confidence intervals for 17 variables used as predictors in presence/absence (A) and abundance (B) species distribution models for Long-billed Curlew across the Northern Great Plains during 2009-2014.

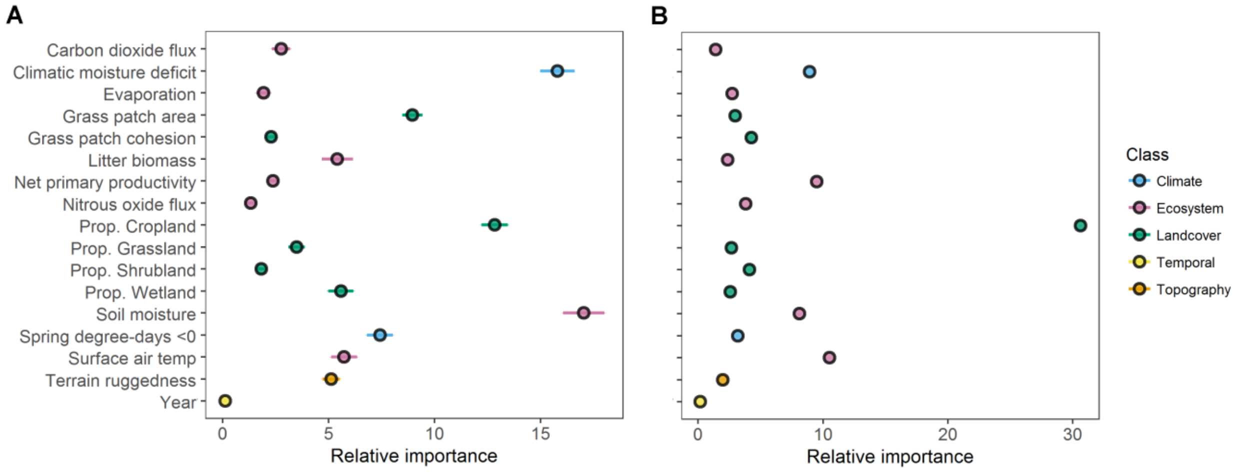

**FIGURE S21.** Mean variable importance and 95% confidence intervals for 17 variables used as predictors in presence/absence (A) and abundance (B) species distribution models for McCown's Longspur across the Northern Great Plains during 2009-2014.

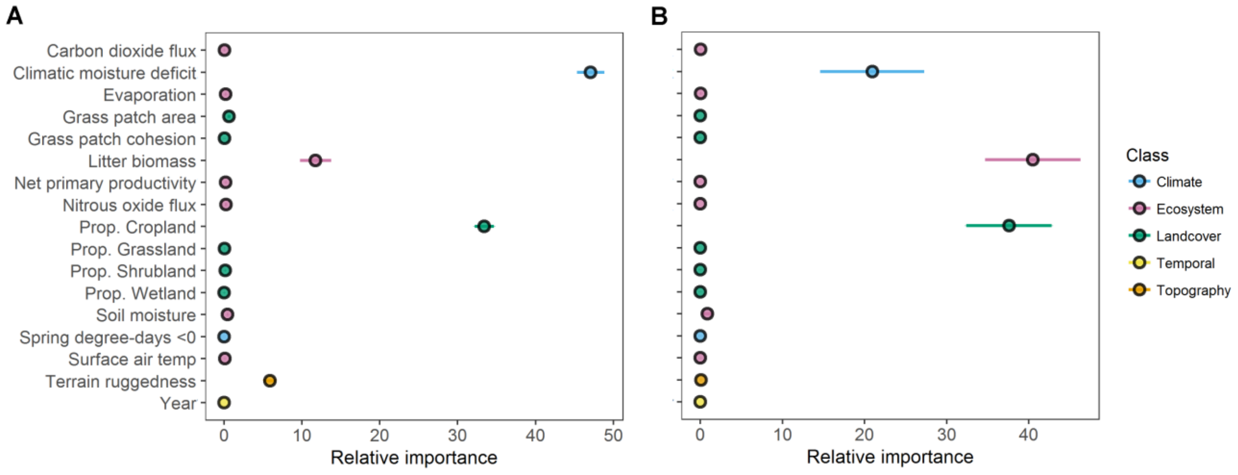


**FIGURE S22.** Mean variable importance and 95% confidence intervals for 17 variables used as predictors in presence/absence (A) and abundance (B) species distribution models for Mountain Plover across the Northern Great Plains during 2009-2014.

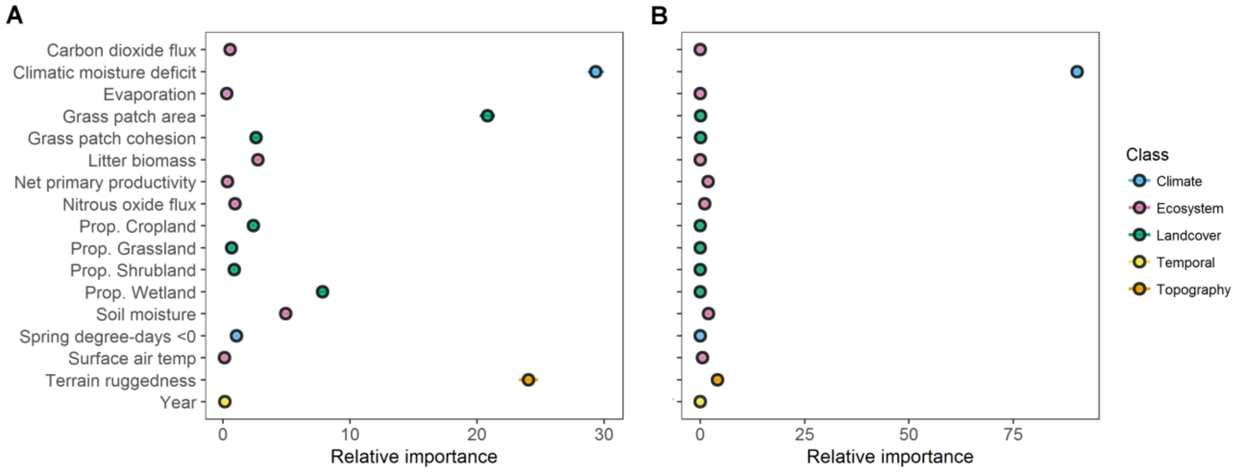

**FIGURE S23.** Mean variable importance and 95% confidence intervals for 17 variables used as predictors in presence/absence (A) and abundance (B) species distribution models for Northern Harrier across the Northern Great Plains during 2009-2014.

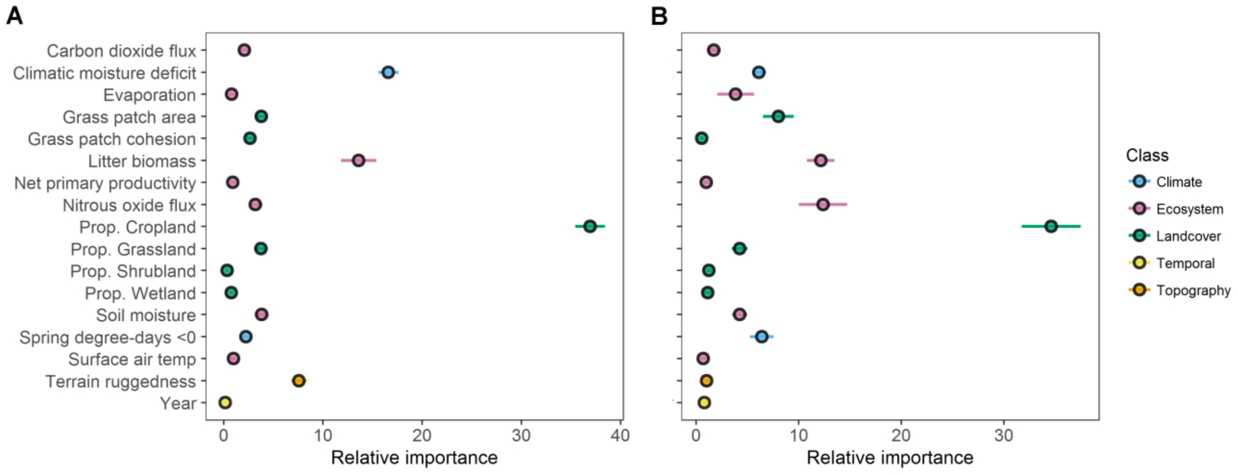


**FIGURE S24.** Mean variable importance and 95% confidence intervals for 17 variables used as predictors in presence/absence (A) and abundance (B) species distribution models for Rock Wren across the Northern Great Plains during 2009-2014.

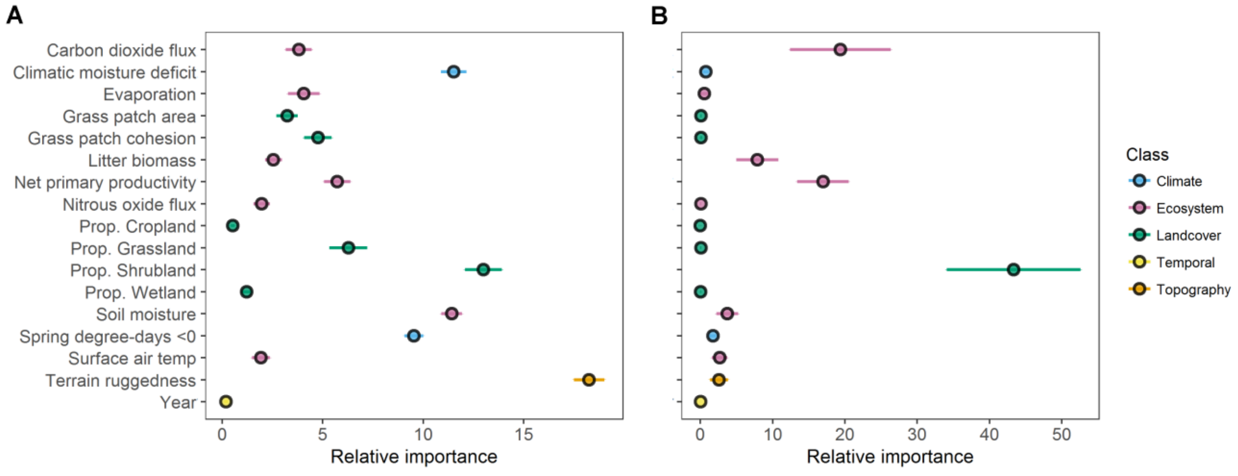

**FIGURE S25.** Mean variable importance and 95% confidence intervals for 17 variables used as predictors in presence/absence (A) and abundance (B) species distribution models for Sage Thrasher across the Northern Great Plains during 2009-2014.

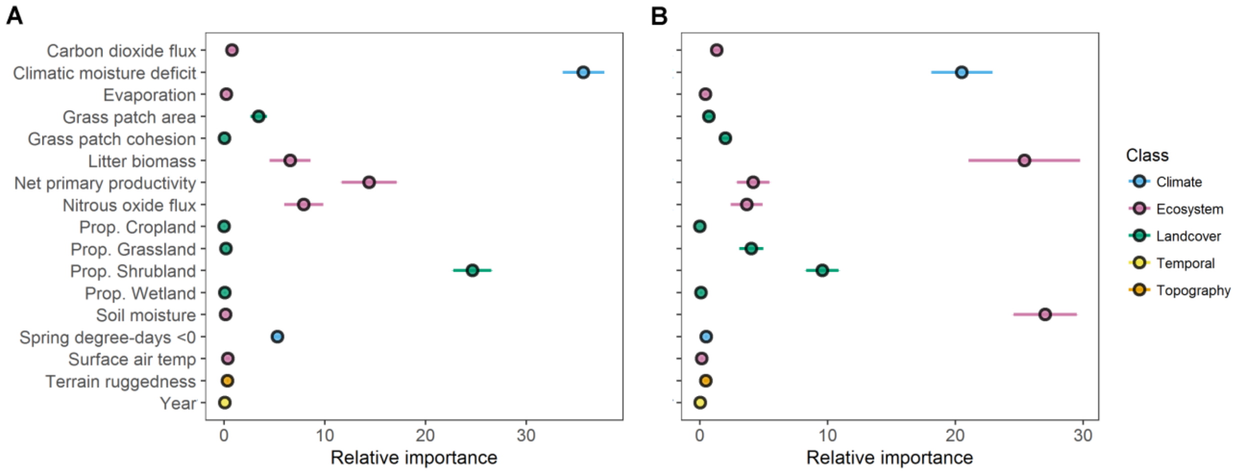


**FIGURE S26.** Mean variable importance and 95% confidence intervals for 17 variables used as predictors in presence/absence (A) and abundance (B) species distribution models for Savannah Sparrow across the Northern Great Plains during 2009-2014.

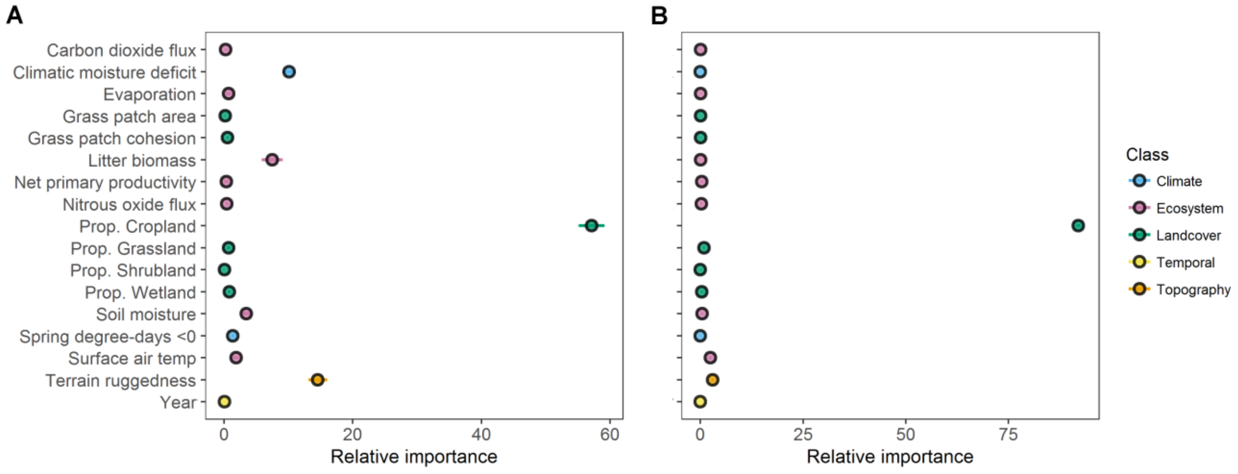

**FIGURE S27.** Mean variable importance and 95% confidence intervals for 17 variables used as predictors in presence/absence (A) and abundance (B) species distribution models for Sharp-tailed Grouse across the Northern Great Plains during 2009-2014.

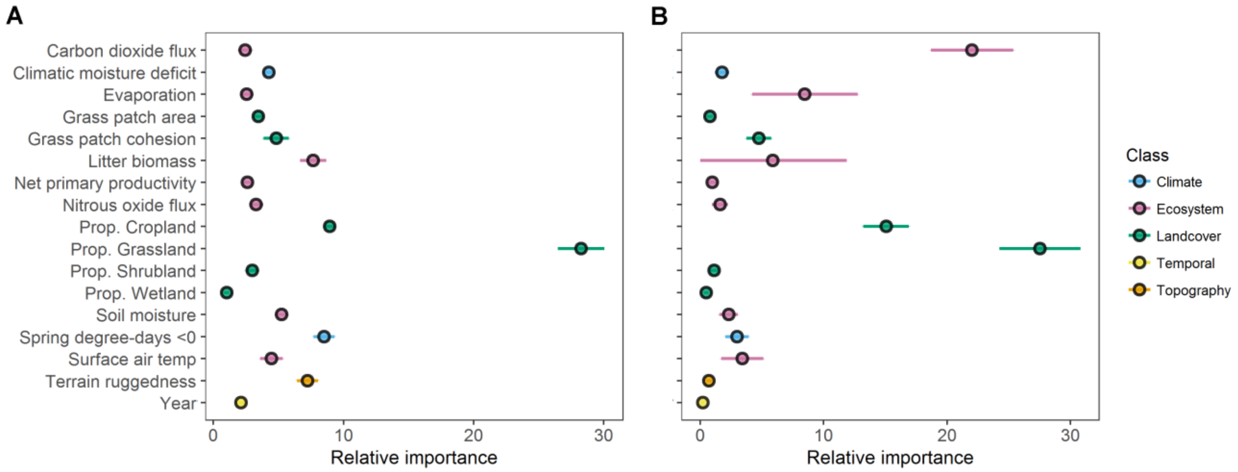


**FIGURE S28.** Mean variable importance and 95% confidence intervals for 17 variables used as predictors in presence/absence (A) and abundance (B) species distribution models for Sprague's Pipit across the Northern Great Plains during 2009-2014.

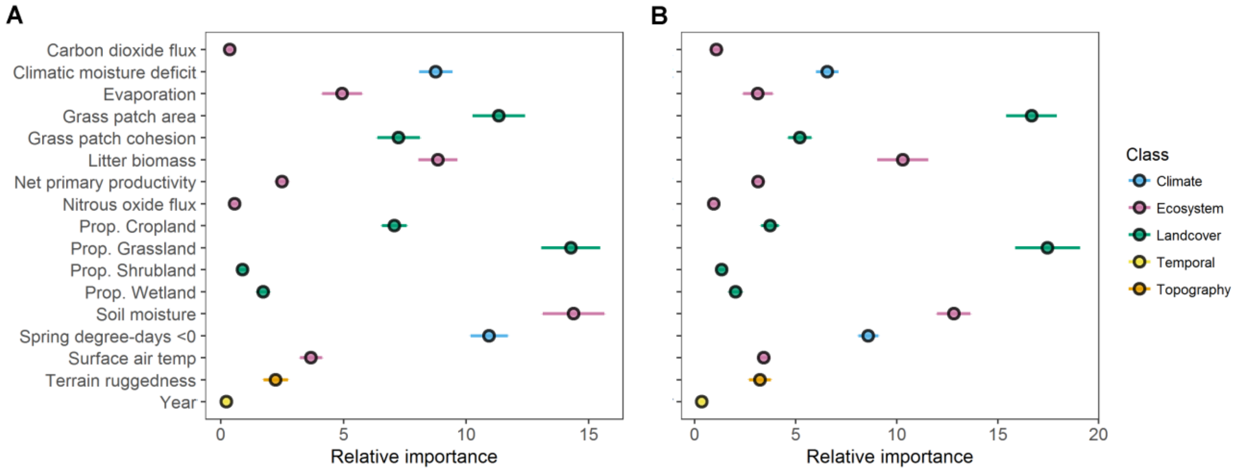

**FIGURE S29.** Mean variable importance and 95% confidence intervals for 17 variables used as predictors in presence/absence (A) and abundance (B) species distribution models for Swainson's Hawk across the Northern Great Plains during 2009-2014.

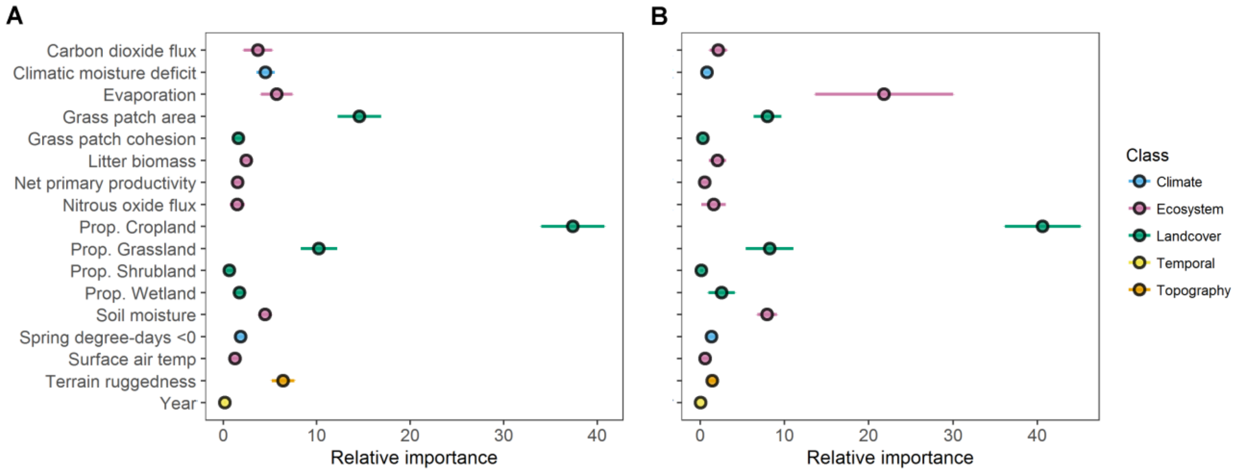


**FIGURE S30.** Mean variable importance and 95% confidence intervals for 17 variables used as predictors in presence/absence (A) and abundance (B) species distribution models for Upland Sandpiper across the Northern Great Plains during 2009-2014.

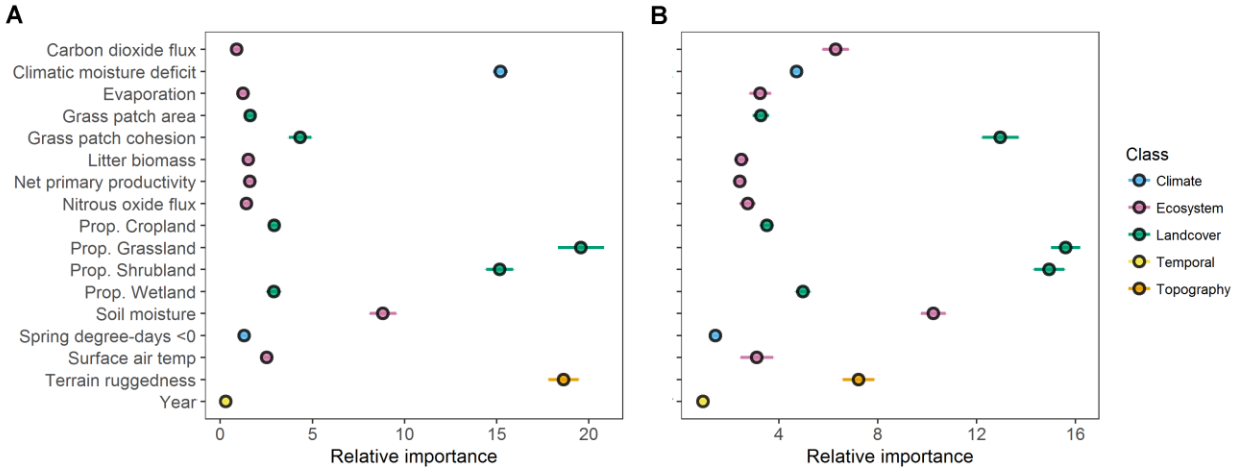

**FIGURE S31.** Mean variable importance and 95% confidence intervals for 17 variables used as predictors in presence/absence (A) and abundance (B) species distribution models for Vesper Sparrow across the Northern Great Plains during 2009-2014.

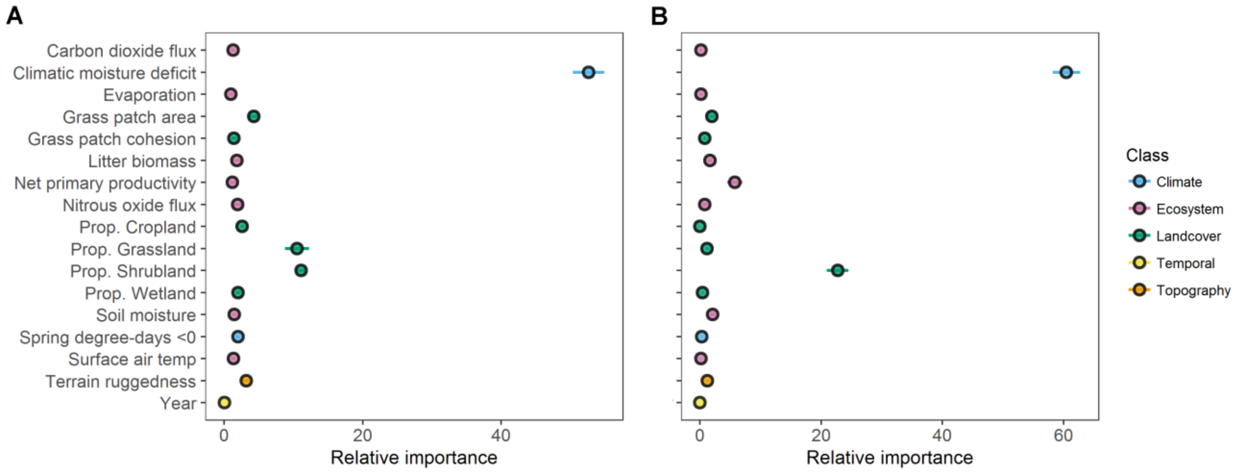


**FIGURE S32.** Mean variable importance and 95% confidence intervals for 17 variables used as predictors in presence/absence (A) and abundance (B) species distribution models for Western Kingbird across the Northern Great Plains during 2009-2014.

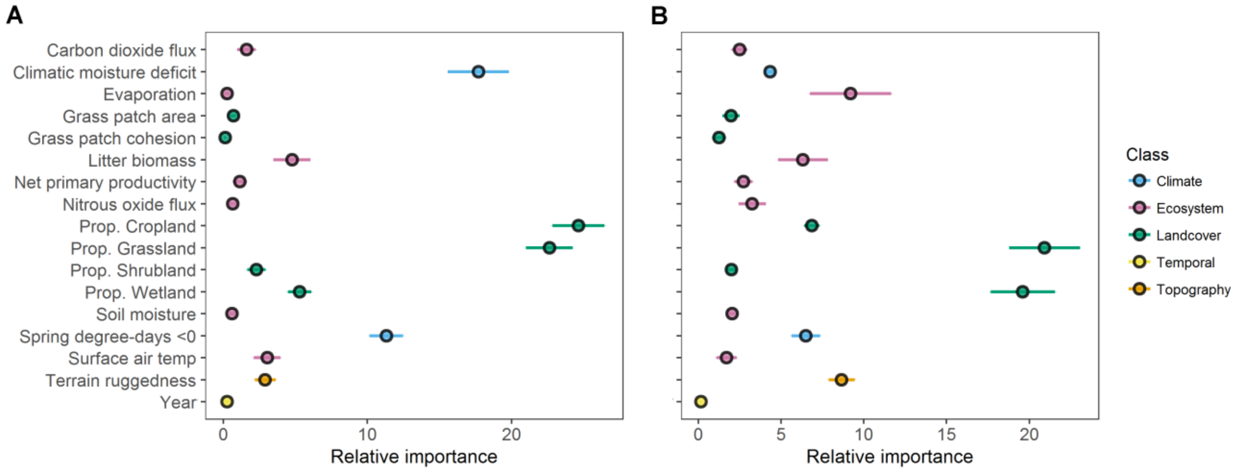

**FIGURE S33.** Mean variable importance and 95% confidence intervals for 17 variables used as predictors in presence/absence (A) and abundance (B) species distribution models for Western Meadowlark across the Northern Great Plains during 2009-2014.

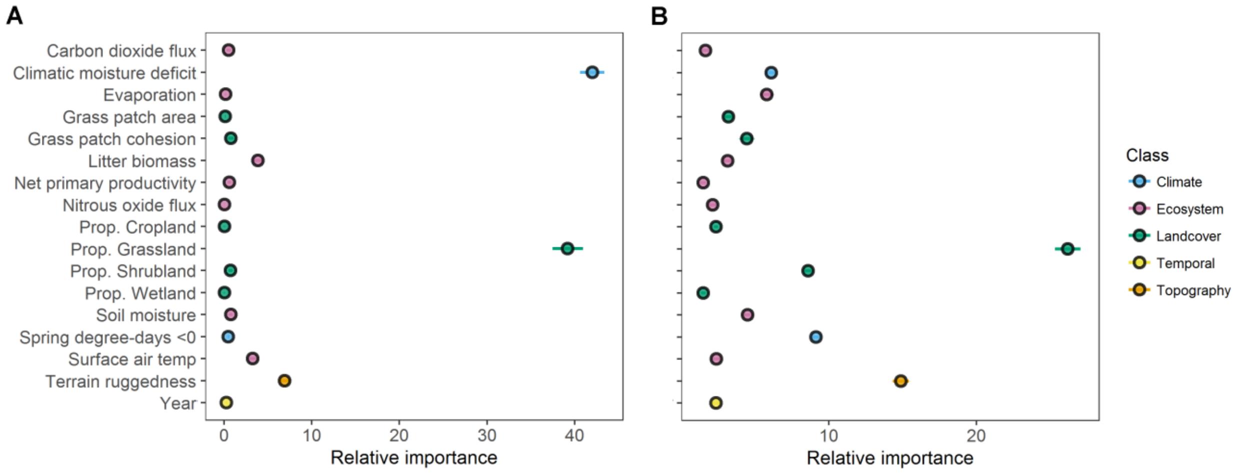


**FIGURE S34.** Mean variable importance and 95% confidence intervals for 17 variables used as predictors in presence/absence (A) and abundance (B) species distribution models for White-throated Swift across the Northern Great Plains during 2009-2014.

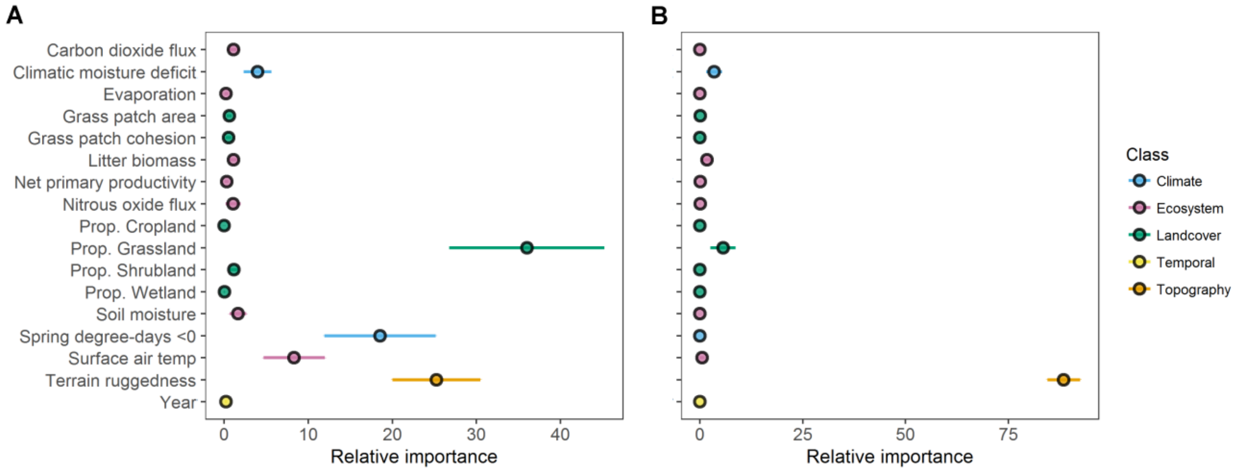
